# Molecular Transducers of Physical Activity Consortium (MoTrPAC): Initial Insights into the Dynamic Human Responses to Exercise

**DOI:** 10.64898/2026.03.02.705347

**Authors:** MoTrPAC Study Group, Anna R Brandt, Jerome Fleg, Bret H Goodpaster, Byron Jaeger, Christopher A Jin, Neil M Johannsen, Daniel Katz, Hasmik Keshishian, Wendy M Kohrt, William E Kraus, Bridget Lester, Edward L Melanson, Michael E Miller, Samuel Montalvo, W Jack Rejeski, Samiya M Shimly, Gregory R Smith, Cynthia L Stowe, Scott Trappe, Hiba AbouAssi, Nicole Adams, David Amar, Euan Ashley, Arianne Aslamy, Marcas M Bamman, Alicia Belangee, Will Bennett, Bryan C Bergman, Daniel H Bessesen, Sue C Bodine, Gerard Boyd, Thomas W Buford, Charles F Burant, Elvis A Carnero, Steven Carr, Toby L Chambers, Clarisa Chavez, Haiying Chen, Shyh-Huei Chen, Jeffrey W Christle, Alex Claiborne, Natalie Clark, Zachary S Clayton, Paul M Coen, Samuel Cohen, Katherine A Collins-Bennett, Dan M Cooper, Tiffany Cortes, Brian Coyne, Gary R Cutter, Matthew Douglass, Shannon S Emilson, Sara Espinoza, Jason Fanning, Will Fountain, Robert E Gerszten, Catherine Gervais, Laurie J Goodyear, Aaron Gouw, Bruce Graham, Kevin Gries, Fadia Haddad, Melissa Harris, Patrick Hart, Andrew Hepler, Joseph A Houmard, Fang-Chi Hsu, Kim M Huffman, Ryan P Hughes, John M Jakicic, Catherine M Jankowski, David Jimenez-Morales, Johanna L Johnson, Raymond Jones, Andrew Jones, Dillon Kuszmaul, Malene E Lindholm, Adam Lowe, Colleen Lynch, Kiril Minchev, Cristhian Montenegro, Kerrie L Moreau, Michael Muehlbauer, Nicolas Musi, Jonathan Myers, Masatoshi Naruse, Christopher B Newgard, Shlomit Radom-Aizik, Tuomo Rankinen, Blake B Rasmussen, Ulrika Raue, Eric Ravussin, Eric W Reynolds, Joseph Rigdon, Ethan Robbins, Jeremy M Robbins, Renee Rogers, Kaitlyn Rogers, Jessica L Rooney, Leanna M Ross, Irene E Schauer, Robert S Schwartz, Stuart C Sealfon, Courtney Simmons, Chad Skiles, Michael P Snyder, Lauren M Sparks, Andrew Stroh, Yifei Sun, Kristen Sutton, Jennifer W Talton, Taylor Taylor, Anna Thalacker-Mercer, Russell Tracy, Todd A Trappe, Caroline S Vincenty, Elena Volpi, Michael P Walkup, Martin J Walsh, Gary Weaver, Matthew T Wheeler, Zhen Yan, Gilhyeon Yoon

**Affiliations:** Ball State University, Muncie, IN; National Institutes of Health, Bethesda, MD; AdventHealth Translational Research Institute for Metabolism and Diabetes, Orlando; Data Management, Analysis, and Quality Control Center - Wake Forest University School of Medicine, Winston-Salem; Stanford University, Stanford, CA; Pennington Biomedical Research Center, Baton Rouge, LA; Bioinformatics Center - Stanford University, Stanford, CA; Broad Institute of MIT & Harvard (Carr Lab), Cambridge, MA; University of Colorado, Denver, CO; Duke University, Durham, NC; Exercise Intervention Core - Wake Forest University, Winston-Salem; Icahn School of Medicine at Mount Sinai, New York City, NY; Cedars-Sinai Medical Center, Los Angeles, California; The University of Alabama at Birmingham - Birmingham, AL; University of Iowa, Iowa City, IA; University of Michigan, Ann Arbor, MI; University of California, Irvine, CA; University of Texas Health Science Center, San Antonio, TX; InstituteIncBroad Institute, Inc., Boston, MA; Joslin Diabetes Center, Boston, MA; East Carolina University, Greenville, NC; University of Pittsburgh, Pittsburgh, PA; Broad (Newgard Lab) - Duke University, Durham, NC; Biospecimens Repository - University of Vermont, Burlington, VT; Virginia Polytechnic Institute and State University, Charlottesville, VA

## Abstract

The goal of the Molecular Transducers of Physical Activity Consortium (MoTrPAC) is to examine the physiological and molecular basis for health benefits in response to acute and chronic exercise. Prior to COVID-19 suspension, generally healthy, sedentary participants (N=206, 18-74y) were randomized to endurance exercise (N=80), resistance exercise (N=81), or non-exercise control (N=45) interventions. The prescribed vigorous acute endurance and resistance exercise bouts induced physiological and metabolic perturbations relative to resting homeostasis. The supervised chronic (3d/wk, 12wk) endurance or resistance training programs robustly improved several physiological parameters (i.e., VO_2_peak, muscular strength). Temporal biospecimen (blood, muscle, and adipose) collections and processing coupled to the acute exercise bouts were highly successful. In most cases, blood, muscle, and adipose samples achieved over 90% success. Endurance and resistance exercise induced distinct acute and chronic physiological responses, which provide a framework to interrogate the molecular basis for health adaptations to these two popular exercise modalities.

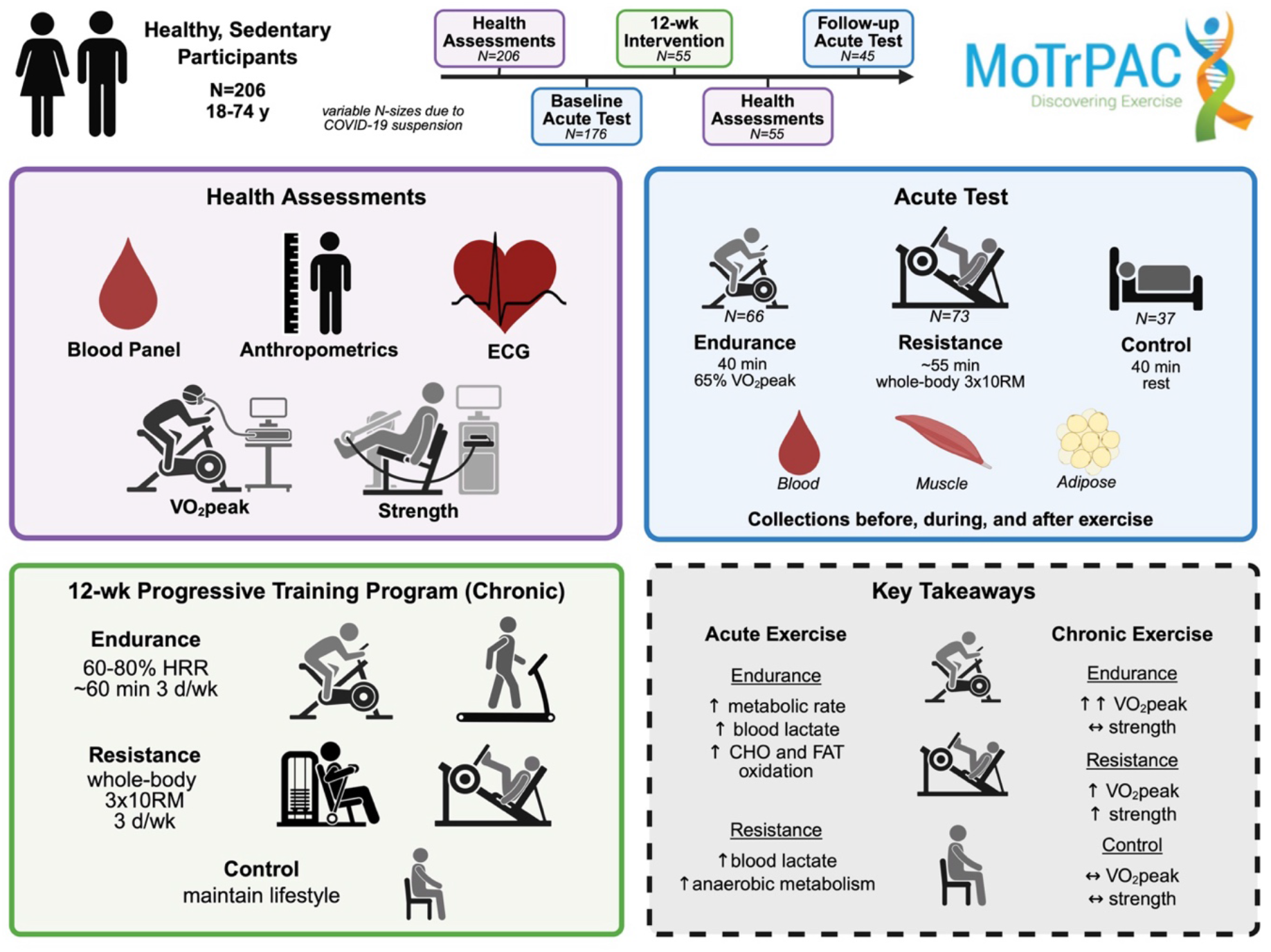

---

Exercise provides a robust physiological stimulus evoking communication (i.e., crosstalk) among multiple tissues and, when repeated regularly (i.e., chronic exercise training), improves physiological capacity, benefits numerous organ systems, and decreases premature morbidity and mortality risk^1–3^. However, a knowledge gap remains in identifying the detailed molecular communication induced by exercise that benefits health and prevents disease^4–6^. The Molecular Transducers of Physical Activity Consortium (MoTrPAC) was established to address these gaps by examining how exercise improves health and mitigates disease by mapping the physiological and molecular responses to acute and chronic exercise^6^. MoTrPAC recently published several original research manuscripts from the pre-clinical rodent studies identifying pronounced physiological improvements [e.g., peak oxygen uptake (VO_2_peak), run speed, and skeletal muscle oxidative capacity] in response to endurance exercise training^7^, which correspond to temporal multi-omic perturbations across multiple organ systems^8–12^. Thousands of shared and tissue-specific molecular alterations were identified across an expansive biological range relevant to numerous aspects of health and disease, with the data available in a public repository (https://motrpac-data.org).

Here, we report initial results from the human randomized trial phase of MoTrPAC representing ∼6-8 months of clinical operations, resulting in 206 sedentary adults randomized into the study. The participants in this report completed various phases of the protocol prior to a temporary cessation of the overall study and closure of this cohort due to the COVID-19 pandemic suspension. This pre-COVID adult human data report aims to: 1) provide initial findings related to the physiological response to endurance and resistance exercise across various phases of the MoTrPAC protocol; 2) highlight study feasibility across several facets of MoTrPAC operations; and 3) discuss lessons learned from this initial human data set as we complete the larger human clinical study. This report accompanies the initial findings on the multi-tissue, multi-omic exercise responses from this same pre-COVID adult human cohort (ref. Molecular Landscape in review at *Nature Medicine*). Further initial interrogation of tissue-specific findings is reported elsewhere ^13–16^.

## RESULTS

### CONSORT Diagram

The CONsolidated Standards Of Reporting Trials (CONSORT) diagram for the sedentary adults recruited prior to suspension of clinical activities due to the COVID-19 pandemic is shown in Figure 1. A total of 206 participants were randomized into three groups: endurance exercise [EE (N=80, 39%)], resistance exercise [RE (N=81, 39%)], and non-exercise control [CON (N=45, 22%)]. By design, a smaller percentage of participants were randomized into CON^17^. A total of 176 participants (85% of those randomized) initiated the pre-intervention baseline acute test (EE, N=66, 83%; RE, N=73, 90%; and CON, N=37, 82%), and among those, 175 (99%) had biospecimen samples collected. Of the 30 participants who did not perform the pre-intervention baseline acute test, 24 (80%) were withdrawn due to the COVID-19 suspension (see Figure 1 legend). Of the 176 participants who initiated the baseline acute test, 145 (82%) began their respective intervention arm; however, only 45 (26%) completed the post-intervention follow-up acute exercise test, with 44 completing the post-intervention biospecimen collections. Of the 121 participants withdrawn after the baseline acute test, 94% were due to the COVID-19 suspension. For a participant to complete all aspects of the study, they visited one of the ten clinical sites 45-48 times for those randomized to exercise (varied slightly due to exercise mode and biospecimens profile, ∼71 hr total) or 15-18 times for CON (∼23 hr total) (Figure 2).

**Figure 1.**
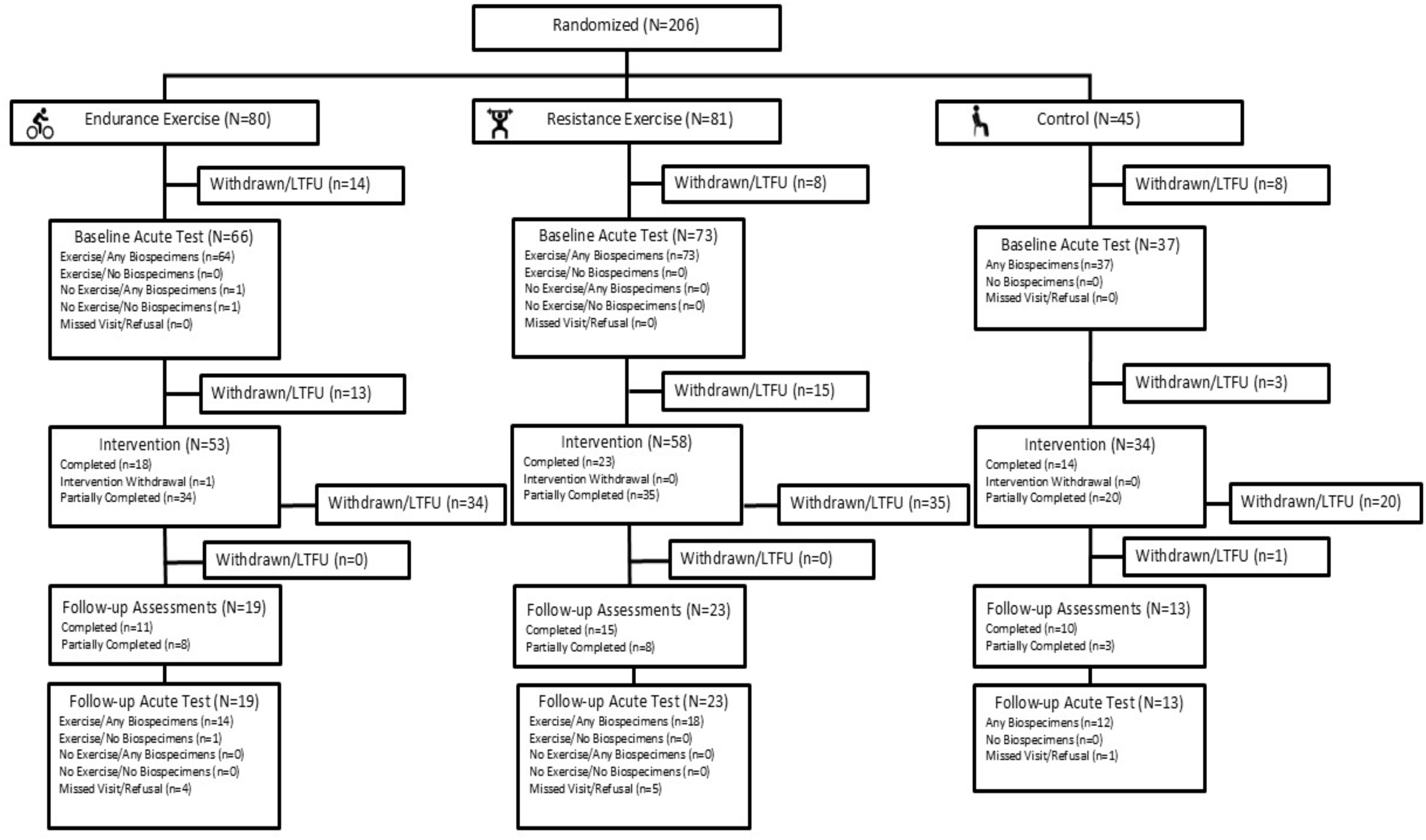
CONSORT diagram. The number of participants eligible for each stage is shown along with the number who completed the stage, partially completed, or withdrew/lost to follow up (LTFU). Participant status in CONSORT diagram allows for incomplete data collection at visits (i.e., only official withdrawals/LFTU are explicitly indicated). Intervention withdrawal indicates that participants were allowed to withdraw from the intervention while remaining in the study for follow-up visits. 91% of participants that were withdrawn/LTFU withdrew due to the COVID-19 suspension; other reasons for participant withdrawal/LTFU included: being too busy (1.3%), conflict with work (1.3%), conflict with caregiving responsibilities (2%), planned travel (0.7%), unwilling to be randomized to control group (0.7%), personal physician opposing involvement (0.7%), study clinician judgement indicating participant is not a good fit (1.3%), unwilling to commit to 3 days per week of training (0.7%). Of the participants randomized, 12% of the participants were withdrawn/LTFU due to COVID-19 suspension before the pre-intervention Baseline Acute Test; 55% were withdrawn/LFTU due to COVID-19 suspension after the Baseline Acute Test.

**Figure 2:**
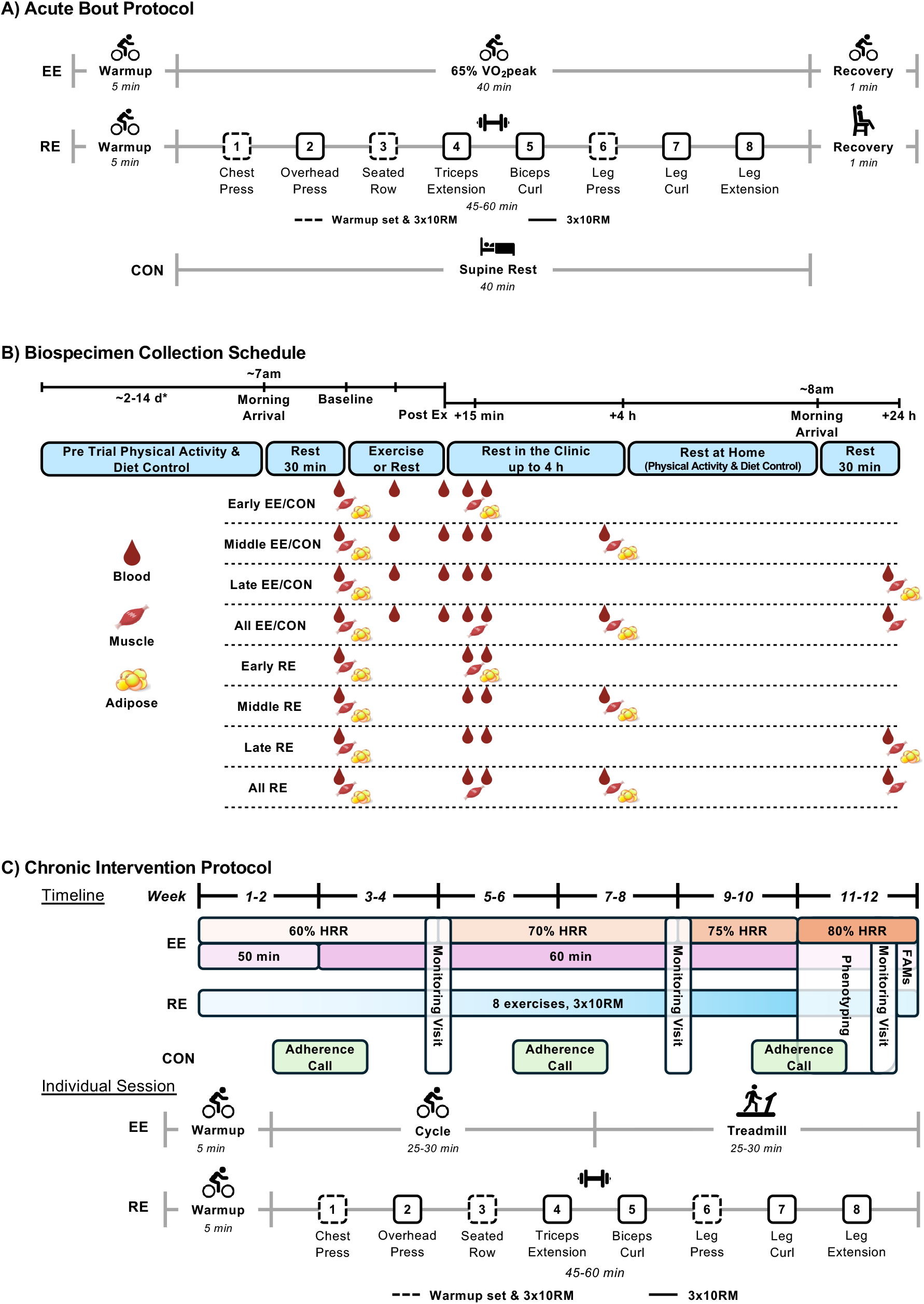
MoTrPAC sedentary adult acute test and chronic intervention protocols. A) Acute bout protocols for endurance, resistance, and control participants. At post-intervention follow-up, only baseline biospecimen samples were collected for CON; no acute rest was completed. B) Acute test biospecimen collection schedule. *Pre trial controls varied at pre-intervention baseline and post-intervention follow-up. C) Overview of protocols for endurance exercise, resistance exercise, and control participants during the 12-week intervention. EE, endurance exercise; RE, resistance exercise; CON, control; VO_2_peak, peak oxygen consumption; 10RM, 10-repetition maximum; Ex, exercise; HRR, heart rate reserve; FAMs, familiarization sessions.

### Baseline Phenotypic Assessments

The pre-screening, medical screening, and baseline phenotyping occurred over 5 visits (∼5 hr total). Baseline demographic and health characteristics for participants who met screening criteria^17^ and were randomized are shown in Table 1. Data are reported as median (25^th^ percentile, 75^th^ percentile) unless stated otherwise. The median age of the cohort was 39 y (range: 18-74 y) and was composed of 72% female participants and 73% White participants (see Table 1 for racial and ethnic breakdown and Supplementary Table 1 for data by sex). Despite dropouts in each study phase due to the COVID-19 suspension, the demographic, health, and phenotypic characteristics at pre-intervention baseline are generally comparable across screening/randomization (N=206; Table 1), baseline acute tests (N=176; see Supplementary Table 2 for characteristics by intervention group; Supplementary Table 3 for characteristics by acute test initiation status), and post-intervention follow-up tests (N=54; Supplementary Table 4).

**Table 1.**
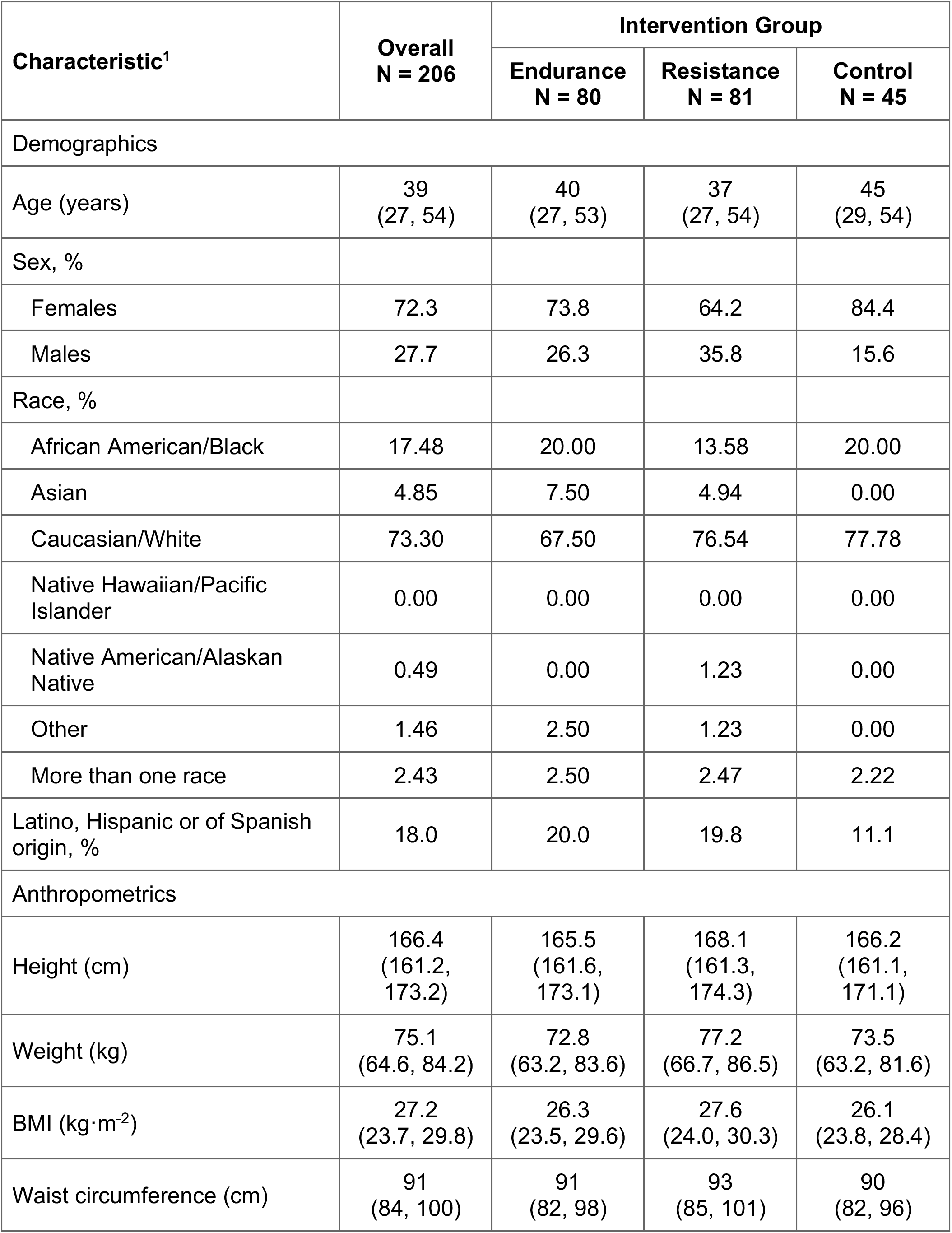
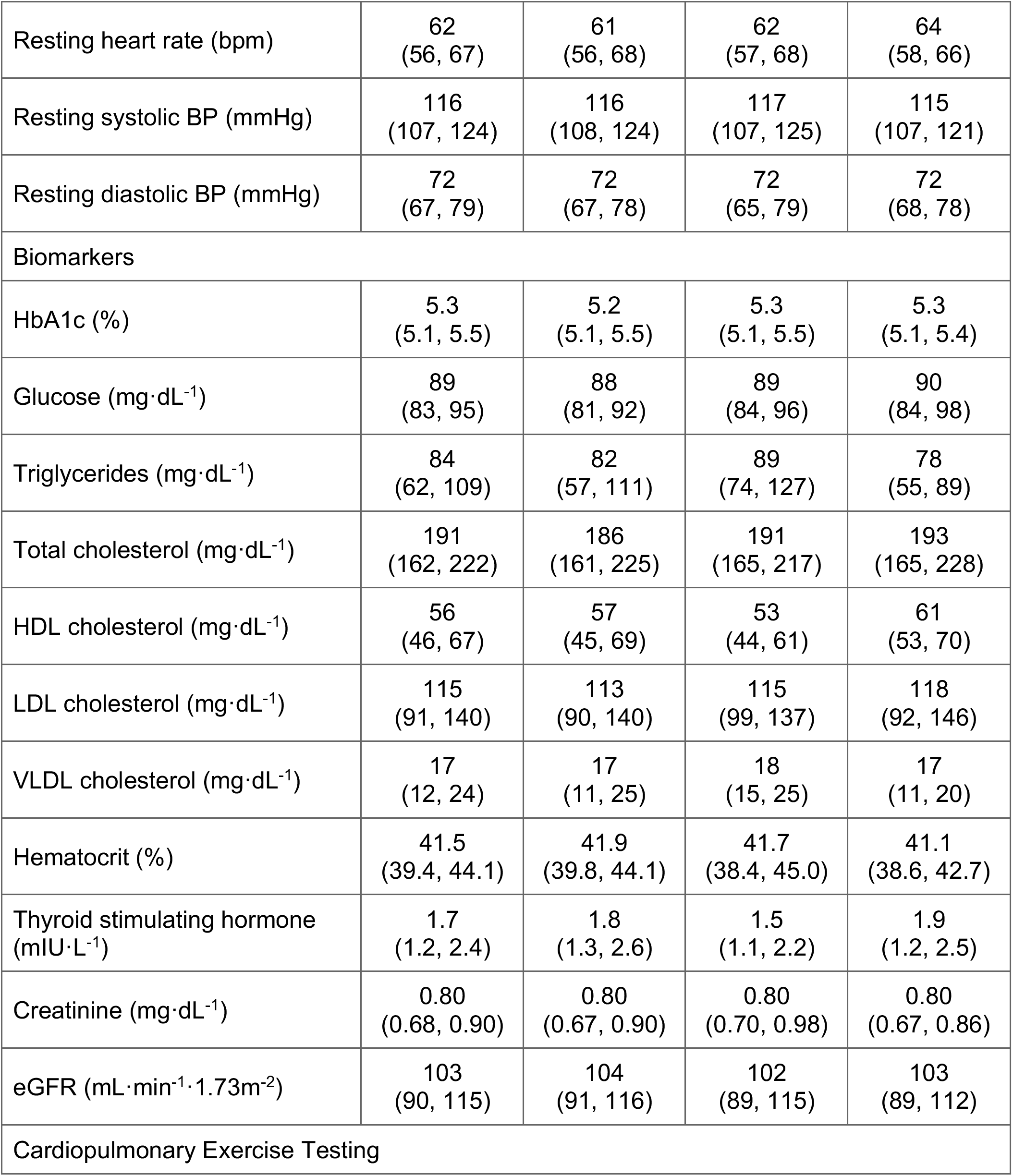
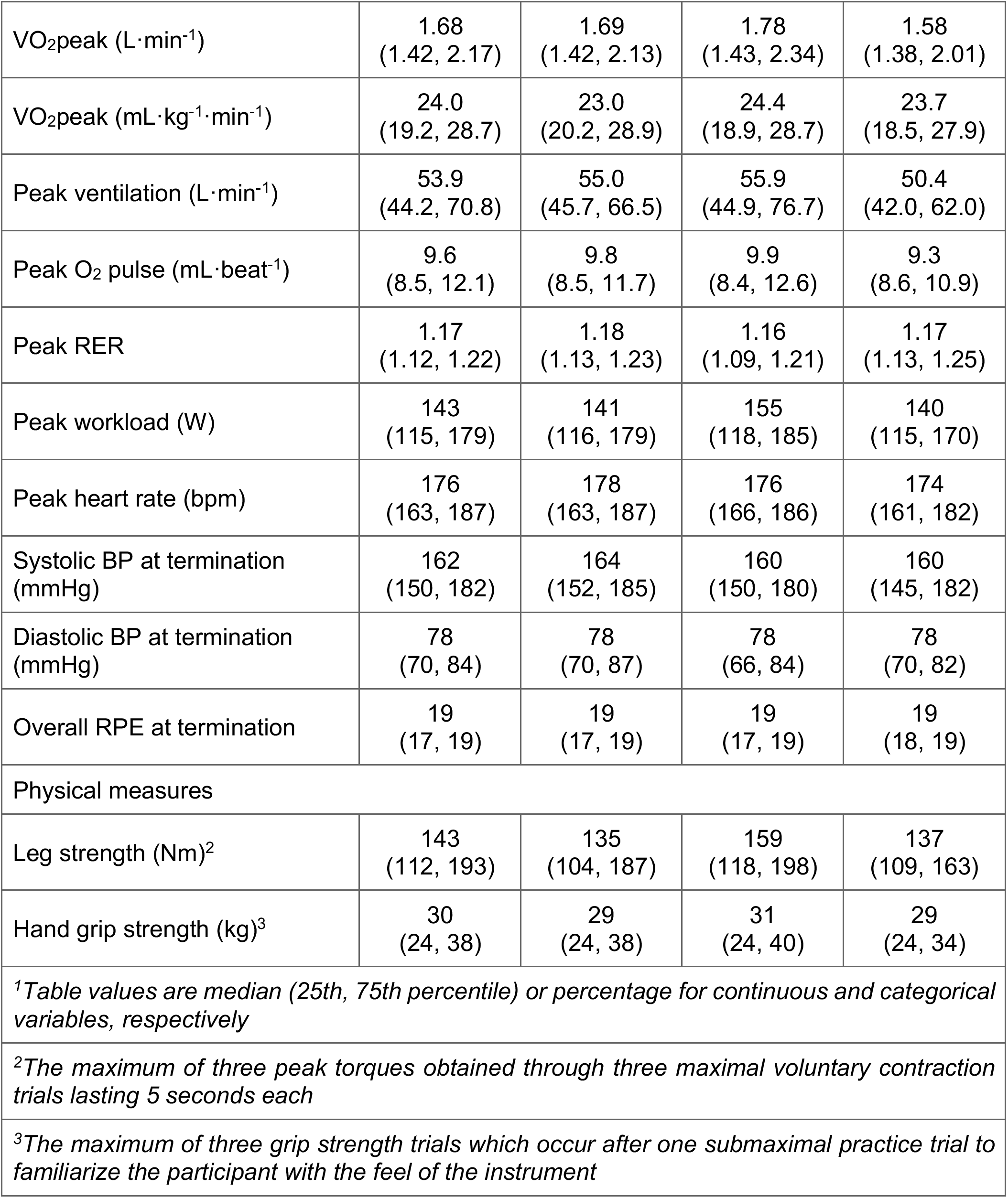
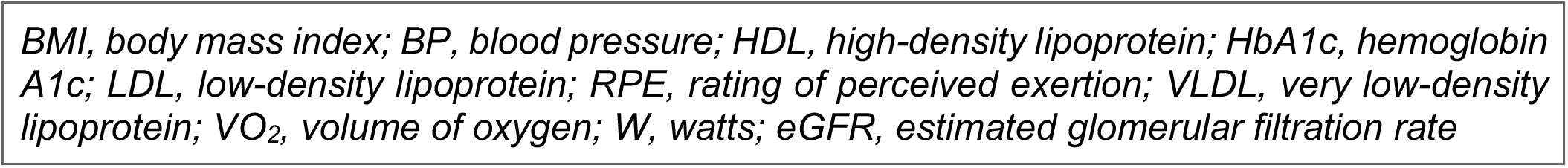
Baseline characteristics of randomized participants by intervention group.

| Characteristic<sup>1</sup> | Overall<br>N = 206 | Intervention Group |  |  |
| --- | --- | --- | --- | --- |
|  |  | Endurance<br>N = 80 | Resistance<br>N = 81 | Control<br>N = 45 |
| Demographics |  |  |  |  |
| Age (years) | 39<br>(27, 54) | 40<br>(27, 53) | 37<br>(27, 54) | 45<br>(29, 54) |
| Sex, % |  |  |  |  |
| Females | 72.3 | 73.8 | 64.2 | 84.4 |
| Males | 27.7 | 26.3 | 35.8 | 15.6 |
| Race, % |  |  |  |  |
| African American/Black | 17.48 | 20.00 | 13.58 | 20.00 |
| Asian | 4.85 | 7.50 | 4.94 | 0.00 |
| Caucasian/White | 73.30 | 67.50 | 76.54 | 77.78 |
| Native Hawaiian/Pacific Islander | 0.00 | 0.00 | 0.00 | 0.00 |
| Native American/Alaskan Native | 0.49 | 0.00 | 1.23 | 0.00 |
| Other | 1.46 | 2.50 | 1.23 | 0.00 |
| More than one race | 2.43 | 2.50 | 2.47 | 2.22 |
| Latino, Hispanic or of Spanish origin, % | 18.0 | 20.0 | 19.8 | 11.1 |
| Anthropometrics |  |  |  |  |
| Height (cm) | 166.4<br>(161.2, 173.2) | 165.5<br>(161.6, 173.1) | 168.1<br>(161.3, 174.3) | 166.2<br>(161.1, 171.1) |
| Weight (kg) | 75.1<br>(64.6, 84.2) | 72.8<br>(63.2, 83.6) | 77.2<br>(66.7, 86.5) | 73.5<br>(63.2, 81.6) |
| BMI (kg·m<sup>-2</sup>) | 27.2<br>(23.7, 29.8) | 26.3<br>(23.5, 29.6) | 27.6<br>(24.0, 30.3) | 26.1<br>(23.8, 28.4) |
| Waist circumference (cm) | 91<br>(84, 100) | 91<br>(82, 98) | 93<br>(85, 101) | 90<br>(82, 96) |
|  |  | Endurance<br>N = 80 | Resistance<br>N = 81 | Control<br>N = 45 |
| Resting heart rate (bpm) | 62<br>(56, 67) | 61<br>(56, 68) | 62<br>(57, 68) | 64<br>(58, 66) |
| Resting systolic BP (mmHg) | 116<br>(107, 124) | 116<br>(108, 124) | 117<br>(107, 125) | 115<br>(107, 121) |
| Resting diastolic BP (mmHg) | 72<br>(67, 79) | 72<br>(67, 78) | 72<br>(65, 79) | 72<br>(68, 78) |
| Biomarkers |  |  |  |  |
| HbA1c (%) | 5.3<br>(5.1, 5.5) | 5.2<br>(5.1, 5.5) | 5.3<br>(5.1, 5.5) | 5.3<br>(5.1, 5.4) |
| Glucose (mg·dL<sup>-1</sup>) | 89<br>(83, 95) | 88<br>(81, 92) | 89<br>(84, 96) | 90<br>(84, 98) |
| Triglycerides (mg·dL<sup>-1</sup>) | 84<br>(62, 109) | 82<br>(57, 111) | 89<br>(74, 127) | 78<br>(55, 89) |
| Total cholesterol (mg·dL<sup>-1</sup>) | 191<br>(162, 222) | 186<br>(161, 225) | 191<br>(165, 217) | 193<br>(165, 228) |
| HDL cholesterol (mg·dL<sup>-1</sup>) | 56<br>(46, 67) | 57<br>(45, 69) | 53<br>(44, 61) | 61<br>(53, 70) |
| LDL cholesterol (mg·dL<sup>-1</sup>) | 115<br>(91, 140) | 113<br>(90, 140) | 115<br>(99, 137) | 118<br>(92, 146) |
| VLDL cholesterol (mg·dL<sup>-1</sup>) | 17<br>(12, 24) | 17<br>(11, 25) | 18<br>(15, 25) | 17<br>(11, 20) |
| Hematocrit (%) | 41.5<br>(39.4, 44.1) | 41.9<br>(39.8, 44.1) | 41.7<br>(38.4, 45.0) | 41.1<br>(38.6, 42.7) |
| Thyroid stimulating hormone<br>(mIU·L<sup>-1</sup>) | 1.7<br>(1.2, 2.4) | 1.8<br>(1.3, 2.6) | 1.5<br>(1.1, 2.2) | 1.9<br>(1.2, 2.5) |
| Creatinine (mg·dL<sup>-1</sup>) | 0.80<br>(0.68, 0.90) | 0.80<br>(0.67, 0.90) | 0.80<br>(0.70, 0.98) | 0.80<br>(0.67, 0.86) |
| eGFR (mL·min<sup>-1</sup>·1.73m<sup>-2</sup>) | 103<br>(90, 115) | 104<br>(91, 116) | 102<br>(89, 115) | 103<br>(89, 112) |
| Cardiopulmonary Exercise Testing |  |  |  |  |
|  |  | Endurance<br>N = 80 | Resistance<br>N = 81 | Control<br>N = 45 |
| VO<sub>2</sub>peak (L·min<sup>-1</sup>) | 1.68<br>(1.42, 2.17) | 1.69<br>(1.42, 2.13) | 1.78<br>(1.43, 2.34) | 1.58<br>(1.38, 2.01) |
| VO<sub>2</sub>peak (mL·kg<sup>-1</sup>·min<sup>-1</sup>) | 24.0<br>(19.2, 28.7) | 23.0<br>(20.2, 28.9) | 24.4<br>(18.9, 28.7) | 23.7<br>(18.5, 27.9) |
| Peak ventilation (L·min<sup>-1</sup>) | 53.9<br>(44.2, 70.8) | 55.0<br>(45.7, 66.5) | 55.9<br>(44.9, 76.7) | 50.4<br>(42.0, 62.0) |
| Peak O<sub>2</sub> pulse (mL·beat<sup>-1</sup>) | 9.6<br>(8.5, 12.1) | 9.8<br>(8.5, 11.7) | 9.9<br>(8.4, 12.6) | 9.3<br>(8.6, 10.9) |
| Peak RER | 1.17<br>(1.12, 1.22) | 1.18<br>(1.13, 1.23) | 1.16<br>(1.09, 1.21) | 1.17<br>(1.13, 1.25) |
| Peak workload (W) | 143<br>(115, 179) | 141<br>(116, 179) | 155<br>(118, 185) | 140<br>(115, 170) |
| Peak heart rate (bpm) | 176<br>(163, 187) | 178<br>(163, 187) | 176<br>(166, 186) | 174<br>(161, 182) |
| Systolic BP at termination (mmHg) | 162<br>(150, 182) | 164<br>(152, 185) | 160<br>(150, 180) | 160<br>(145, 182) |
| Diastolic BP at termination (mmHg) | 78<br>(70, 84) | 78<br>(70, 87) | 78<br>(66, 84) | 78<br>(70, 82) |
| Overall RPE at termination | 19<br>(17, 19) | 19<br>(17, 19) | 19<br>(17, 19) | 19<br>(18, 19) |
| Physical measures |  |  |  |  |
| Leg strength (Nm)<sup>2</sup> | 143<br>(112, 193) | 135<br>(104, 187) | 159<br>(118, 198) | 137<br>(109, 163) |
| Hand grip strength (kg)<sup>3</sup> | 30<br>(24, 38) | 29<br>(24, 38) | 31<br>(24, 40) | 29<br>(24, 34) |
| <sup>1</sup>Table values are median (25th, 75th percentile) or percentage for continuous and categorical variables, respectively |  |  |  |  |
| <sup>2</sup>The maximum of three peak torques obtained through three maximal voluntary contraction trials lasting 5 seconds each |  |  |  |  |
| <sup>3</sup>The maximum of three grip strength trials which occur after one submaximal practice trial to familiarize the participant with the feel of the instrument |  |  |  |  |
|  |  | Endurance<br>N = 80 | Resistance<br>N = 81 | Control<br>N = 45 |
| *BMI, body mass index; BP, blood pressure; HDL, high-density lipoprotein; HbA1c, hemoglobin A1c; LDL, low-density lipoprotein; RPE, rating of perceived exertion; VLDL, very low-density lipoprotein; VO<sub>2</sub>, volume of oxygen; W, watts; eGFR, estimated glomerular filtration rate* |  |  |  |  |

Cardiorespiratory values obtained at baseline are presented in Table 1 with individual values for males and females plotted across the adult lifespan in Figure 3A. VO_2_peak was 2.67 (2.19, 3.03) L·min^−1^ in males and 1.54 (1.34, 1.84) L·min^−1^ in females. Peak respiratory exchange ratio (RER) was 1.17 (1.12, 1.22), with 94% of participants above a RER of 1.05 (Table 1). RPE at test termination was 19 (17,19) on the Borg scale (6-20)^18^. Peak values obtained for ventilation [VE (L·min^−1^)], O_2_ pulse (a surrogate for stroke volume, ml•beat^−1^), heart rate [HR (bpm)], blood pressure [BP (mmHg)], and workload (watts) are also presented by randomization group in Table 1, and by sex in Supplementary Table 1.

**Figure 3.**
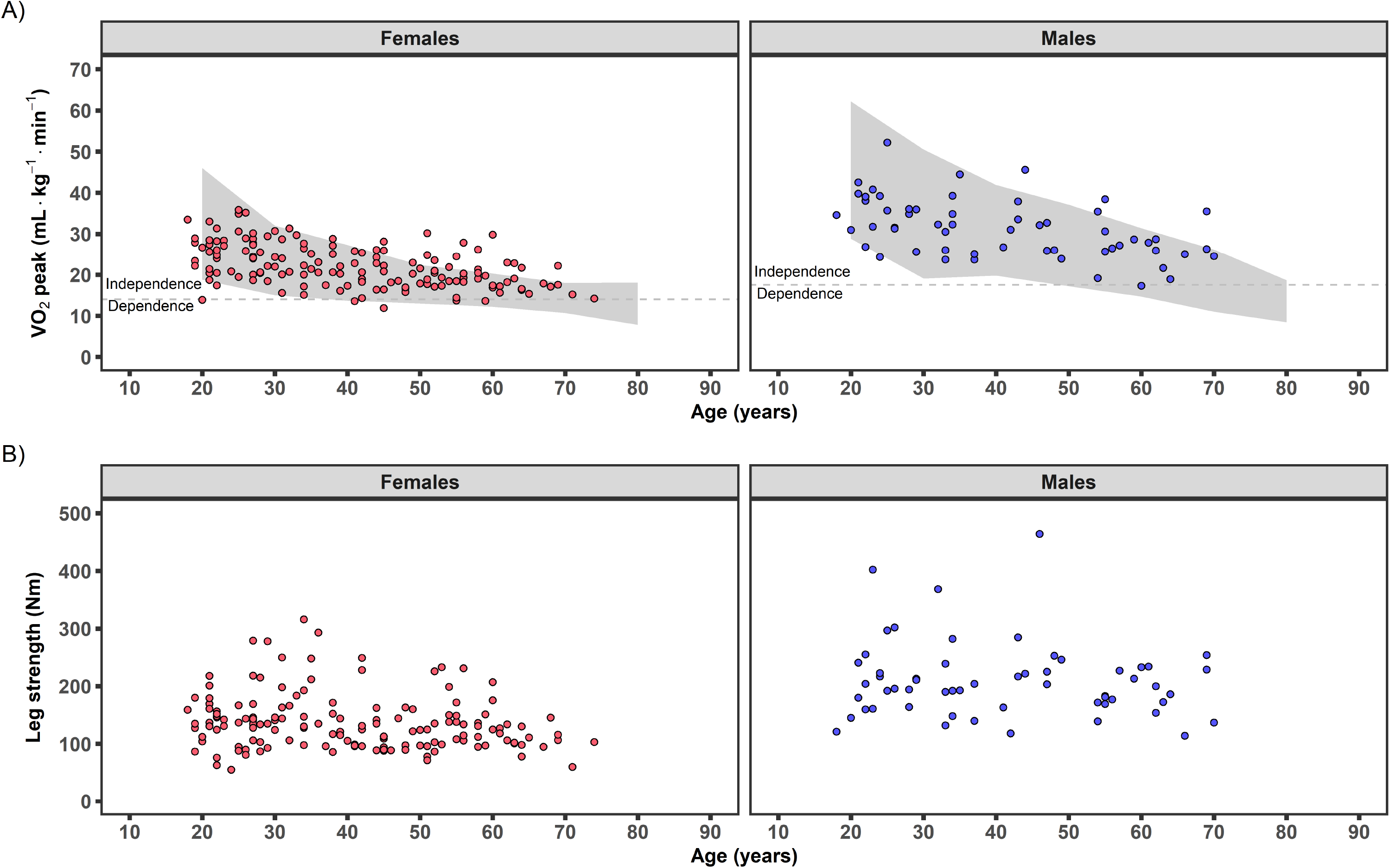
Individual cardiorespiratory fitness and leg strength by age. A) Individual relative VO_2_peak data at pre-intervention baseline for randomized participants (N=206; 149F, 57M) by age compared with normative values from the FRIEND database (3662 females, 2439 males)^20^. Due to the similar exercise test mode (cycle) and direct gas analysis, the FRIEND database was chosen for comparison. Grey area represents the 10^th^-90^th^ percentiles of the FRIEND database across the adult lifespan. The estimated frailty threshold for independence is shown (dotted line) and failure to remain above this threshold may result in increased risk for mortality^3^. B) Individual maximal leg strength (isometric knee extensor strength) for females (N=149) and males (N=57) by age. FRIEND, Fitness Registry and the Importance of Exercise National Database; VO_2_peak, peak oxygen consumption.

Isometric knee extensor strength obtained at baseline is presented in Table 1 with individual values for males and females across the adult lifespan plotted in Figure 3B. Isometric knee extensor strength was 200 (169, 233) Nm in males and 131 (102, 152) Nm in females. Handgrip values are presented in Table 1 with individual values for males and females plotted across the adult lifespan in Ext. Data Figure 1A. Handgrip strength was 45 (38, 50) kg in males and 26 (22, 31) kg in females.

Descriptive profiles (see Supplementary Table 1 for sex and age breakdown) suggest females had higher resting HR, lower absolute (L•min^−1^) and relative (ml•kg^−1^•min^−1^) VO_2_peak, and lower isometric knee extensor muscle strength compared to males. Compared to younger adults (18-39 y), middle-aged and older adults (40+ y) had lower absolute (L•min^−1^) and relative (ml•kg^−1^•min^−1^) VO_2_peak, as well as lower peak HR and higher BP in the final minutes of the cardiopulmonary exercise test (CPET).

Baseline screening interview questionnaires assessing regular physical activity patterns and frequency classified participants as sedentary. Sedentary behavior was also assessed via collection of daily step counts at baseline using wrist-worn actigraphy for 193 participants (140F, 53M) with at least five compliant (≥12 hr wear time) days of data. Average daily step count was below 5000 steps for most participants (81%) with individual values plotted across age in Ext. Data Figure 1B.

### Pre-Intervention Baseline Acute Exercise Bout

Exercise intensity across the EE and RE familiarization sessions and acute bouts was compared to target intensity in Ext. Data Figure 2. The percentage of EE participants falling within the goal of 65±5% VO_2_peak improved from 26% at familiarization session 1 to 83% during the acute bout. The percentage of RE participants falling within the goal of 8-12 repetitions also improved from 59% at familiarization session 2 to 94% during the acute bout. The EE, RE, or CON acute bout with biospecimen sampling resulted in acute tests taking 3.5-6 hr to execute, depending upon the biospecimen temporal profile.

### Baseline Acute Endurance Exercise

Sixty-four participants (46F) performed the 40 min baseline EE acute bout at 64.8% VO_2_peak (63.7, 65.9%; Figure 4A), matching the target intensity of 65±5% VO_2_peak^17^. Absolute VO_2_ of 1.17 L·min^−1^ (1.08, 1.28 L·min^−1^; Figure 4B) was obtained with a workload of 61 W (47, 88 W; Figure 4C) and a total work output of 141 KJ (110, 210 KJ; Supplementary Table 5). This exercise intensity elicited a RER of 0.92 (0.90, 0.93) resulting in 72% (68, 76%) carbohydrate utilization during the exercise bout (Figure 4D), HR of 141 bpm (137, 144 bpm; Figure 4E), and a heart rate reserve (HRR) of 70% (67, 72%). Blood lactate was 3.7 mM (3.2, 4.9 mM) at 20 min of exercise and increased slightly by the end of exercise [40 min: 4.0 mM (2.7, 5.2 mM); Figure 5A].

**Figure 4.**
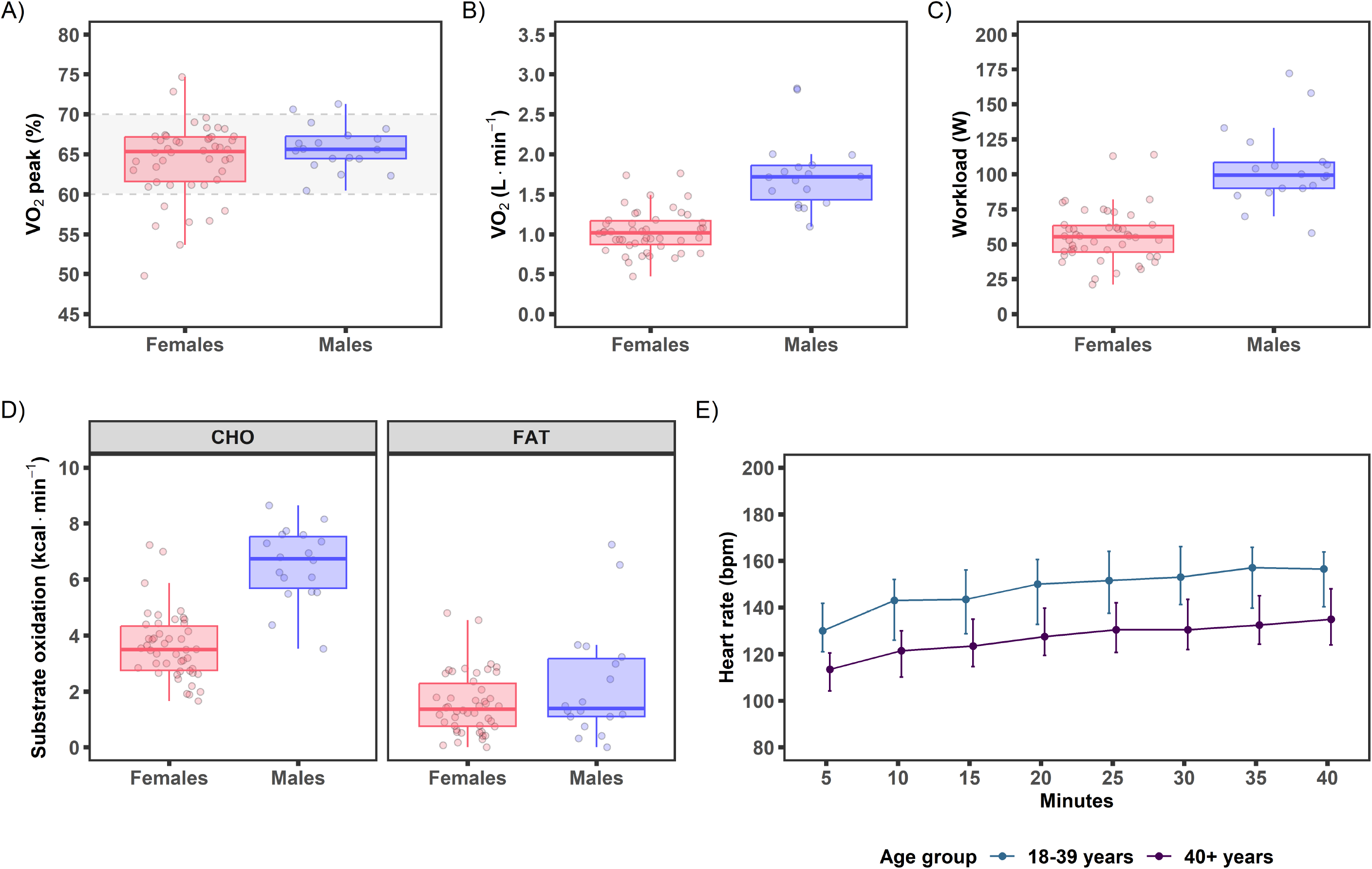
Pre-intervention baseline acute endurance exercise bout parameters shown by sex or age group. A-C) Exercise intensity as (A) percent VO_2_peak (%), (B) VO_2_ (L·min^−1^), and (C) workload (W). Target exercise intensity range of 65±5% VO_2_peak is indicated by the dotted lines and grey shading. D) Average carbohydrate (CHO) and fat utilization during the acute test. E) Following the 5 min warm-up, the median (25^th^-75^th^ percentile) heart rate progression is plotted during the 40 min acute bout over 5 min intervals. Data shown in panels A-D are boxplots and individual observations. Individual observations in panels A-D are 6 min averages of minutes 14-17 with 34-37.

**Figure 5.**
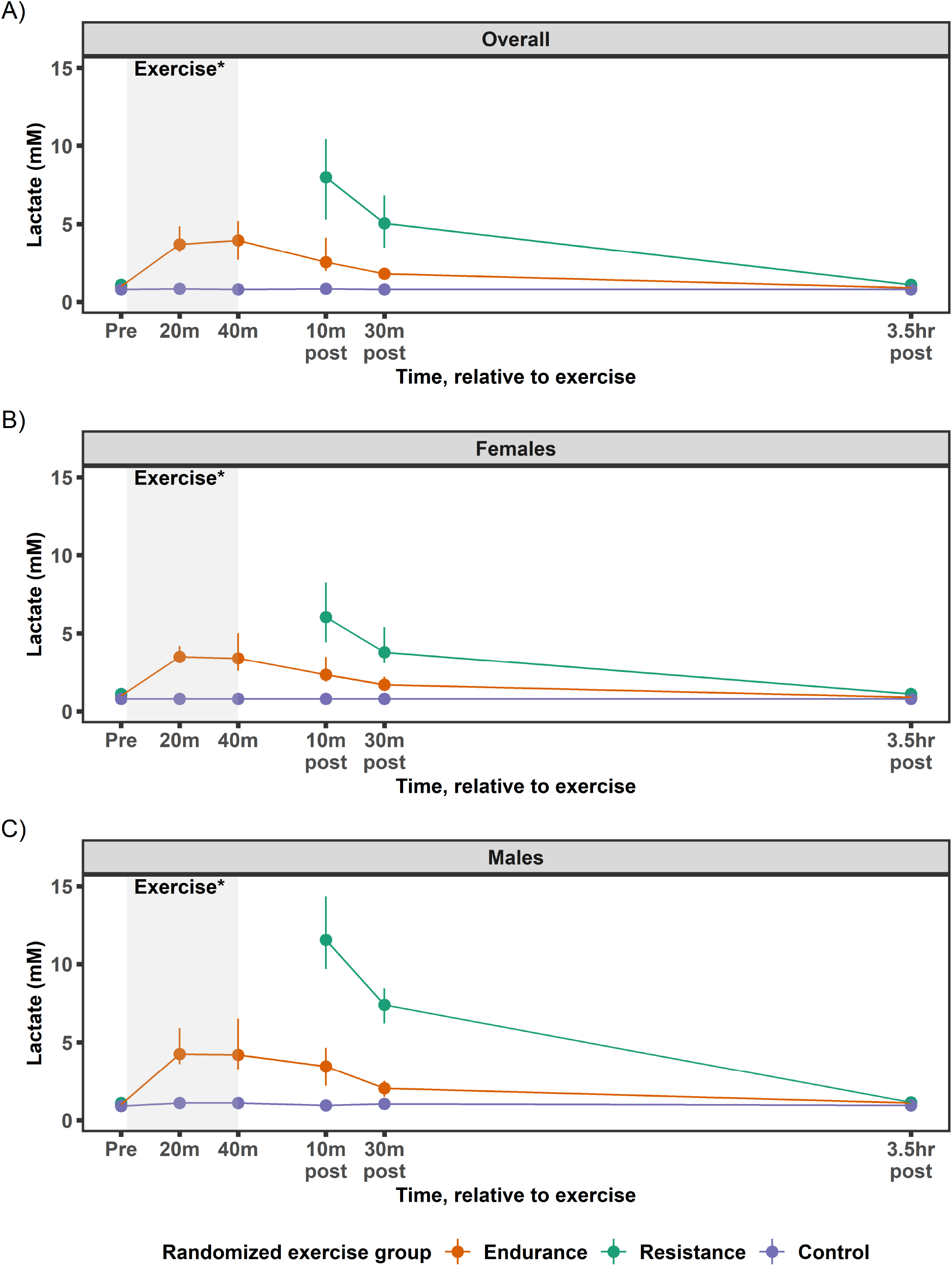
Venous blood lactate from acute endurance, resistance, and control tests. Pre-intervention baseline acute tests shown in A) all participants, B) female participants, and C) male participants for endurance exercise, resistance exercise, and control. Data shown as median (25^th^-75^th^ percentile).

Female and male participants completed the EE acute bout at similar relative intensities (Figure 4A). However, males cycled at a higher workload, which elicited a higher absolute VO_2_, utilized more carbohydrate (relative and absolute), and achieved higher blood lactate during the exercise bout. HR increased during exercise with older participants (40+ y) showing a ∼15-25 bpm lower response over the course of the exercise bout compared to the young (18-39 y).

### Baseline Acute Resistance Exercise

Seventy-three participants (49F) performed the baseline RE acute bout lasting 56 min (51, 59 min; including exercise and rest periods). Participants completed 9.7 (9.3, 10.1) repetitions per set, falling within the target range of 8-12 for 10-repetition maximum (10RM) (Figure 6A). Total loads for upper body, lower body, and combined upper and lower body were 2756 kg (1871, 3785 kg), 4892 kg (3681, 6195 kg), and 7537 kg (5783, 9840 kg), respectively (Figure 6B). Acute bout resistances for chest press, leg press, and leg extension elicited intensities of 62% (54, 71%), 72% (64, 76%), and 59% (51, 64%) of one-repetition maximum (1RM), respectively (Figure 6C). Blood lactate was 8.0 mM (5.3, 10.4 mM) 10 min post-exercise (Figure 5A).

**Figure 6.**
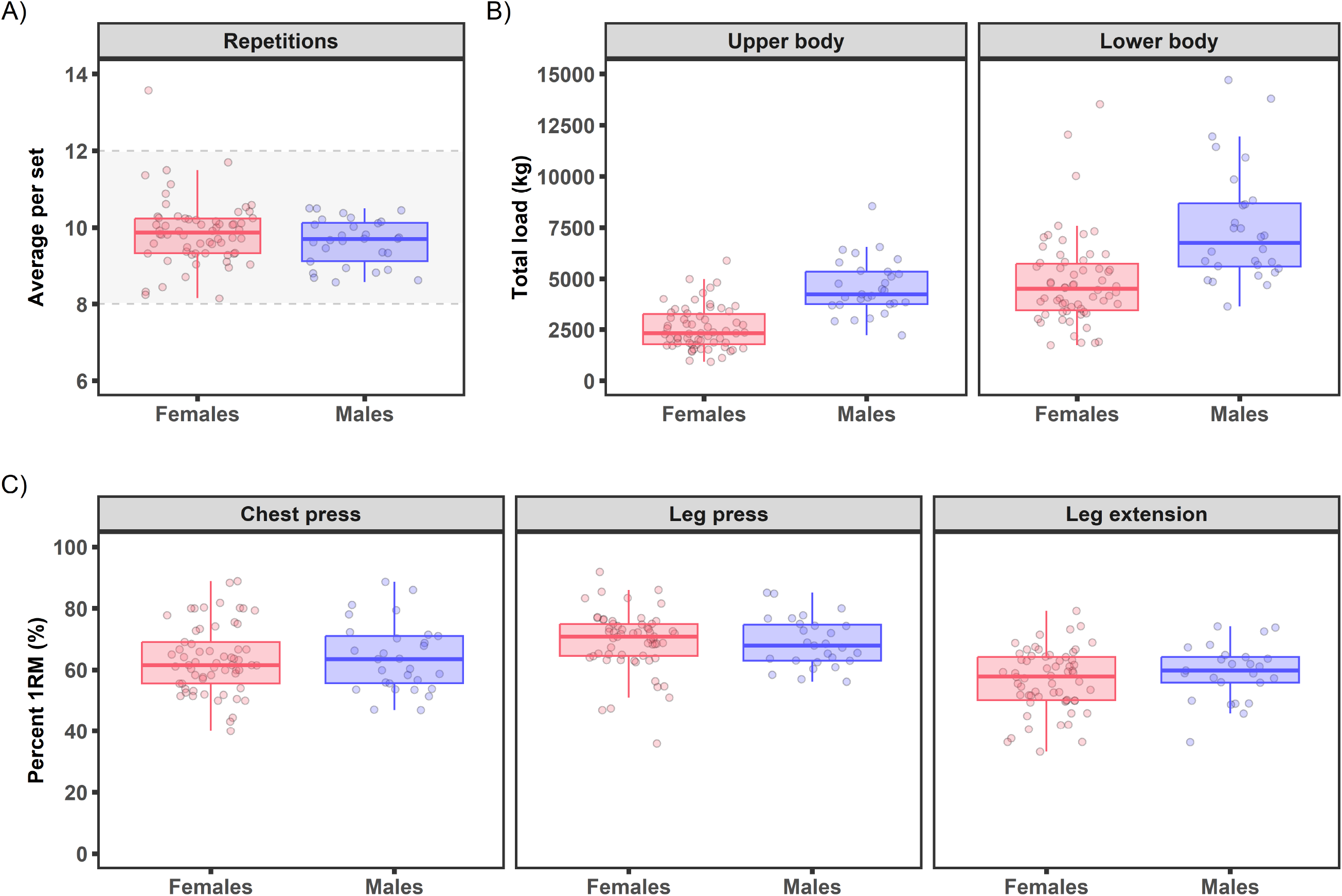
Pre-intervention baseline acute resistance exercise bout parameters by sex. A) Average repetitions per set on all working sets plotted with the 8-12 repetition goal, which is indicated by the dotted lines and grey shading. B) Total load [weight (kg)*repetitions*sets] for lower and upper body. Total load was plotted for participants that completed at least 1 exercise. C) Average percent one repetition maximum (%1RM) for chest press, leg press, and leg extension. Data shown are boxplots with individual observations drawn as points.

Female and male participants completed the RE bout similarly, with comparable average repetitions per set across the eight resistance exercise movements [females: 9.9 (9.3, 10.0); males: 9.7 (9.1, 10.0)] (Figure 6A, see Supplementary Table 6 for sex and age profiles). Likewise, the relative effort as gauged by percent 1RM for chest press, leg press, and leg extension was similar between females and males (Figure 6C). However, the total load was greater in males [11065 kg (9159, 12613 kg)] than females [6228 kg (5305, 7709 kg); Figure 6B, Supplementary Table 6]. Blood lactate increased ∼11-fold in males from rest [1.1 mM (0.9, 1.4 mM)] to 10 min post-exercise [11.6 mM (9.7, 14.4 mM)] and ∼6-fold in females from rest [1.1 mM (0.9, 1.3 mM)] to 10 min post-exercise [6.1 mM (4.4, 8.3 mM); Figure 5B-C]. No notable differences were observed in the resistance exercise stimuli (i.e., percent 1RM, average repetitions) within each sex by age group (18-39 y vs. 40+ y; Supplementary Table 6).

### Intervention and Post-Intervention Follow-Up Phenotypic Assessments

A total of 55 participants (EE=18, RE=23, CON=14) completed their intervention program prior to the COVID-19 suspension (Figure 1). Attendance at the intervention sessions averaged 93% for EE and 92% for RE training groups. The target goals for exercise frequency, intensity, and duration were generally achieved for both intervention groups (Supplementary Table 7). For EE, the median METmin·week^−1^ increased from 677 METmin·week^−1^ during week 2 to 1072 METmin·week^−1^ during week 10, a 47% relative improvement. For RE, the total amount of weight lifted per week increased from 25,055 kg·week^−1^ to 32,002 kg·week^−1^, a 21% relative improvement from week 2 to week 10 (Supplementary Table 7).

Fifty-five participants (EE=19, RE=23, CON=13) participated in post-intervention follow-up phenotypic assessments. The EE and RE groups showed improvements in cardiorespiratory fitness. VO_2_peak (L·min ^−1^) increased 15% (9, 18%) in EE and 8% (1, 14%) in RE. Oxygen pulse increased 17% (11, 20%) in EE and 11% (4, 17%) in RE. Peak workload increased 20% (12, 25%) in EE and 9% (5, 18%) in RE. The CON group was relatively unchanged over the intervention period (Figure 7 and Ext. Data Figure 3, Supplementary Table 8). For strength measures, the RE group improved knee extension strength by 12% (−3, 21%), while pre- to post-intervention knee extension strength was similar in the EE and CON groups (Figure 7 and Ext. Data Figure 3, Supplementary Table 8).

**Figure 7.**
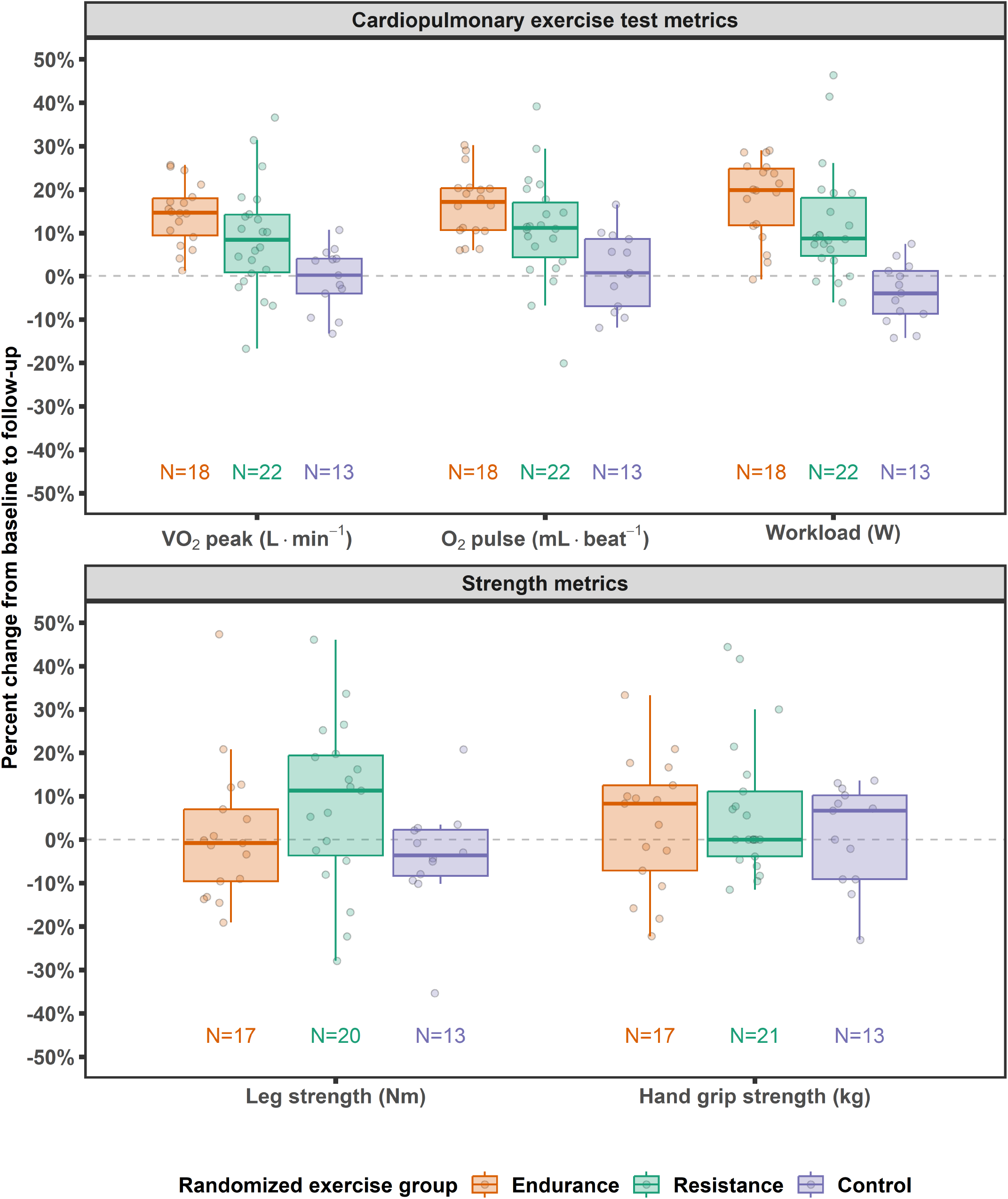
Percent change in maximal CPET and strength metrics from pre-intervention baseline to post-intervention follow-up. Percent change in VO_2_peak (L·min^−1^), peak O_2_ pulse (mL·beat^−1^), and peak workload (W) from the cardiopulmonary exercise test (CPET). Percent change in strength measurements, including maximal leg strength [isometric knee extensor strength (Nm)] and maximal hand grip strength (kg).

### Post-Intervention Follow-up Acute Test

Forty-five participants (EE=15, RE=18, CON=12) completed the post-intervention follow-up acute test. One participant who completed the follow-up EE acute test was not included in the follow-up acute bout data as they opted out of the baseline acute bout.

#### Endurance Exercise

Fourteen participants (11F) completed pre-intervention baseline and post-intervention follow-up EE acute bouts with similar relative exercise intensity, HR, and blood lactate between tests (Ext. Data Figures 4A, 4E, 5A). The workload during the acute bout increased by 30% (24, 82%; Ext. Data Figure 4C), eliciting a 10% (7, 15%) higher absolute VO_2_ (Ext. Data Figure 4B) with corresponding increases in O_2_ pulse [9% (1, 22%)] and VE [21% (9, 43%)] observed after the intervention (Supplementary Table 9). Absolute carbohydrate utilization increased 14% (8, 32%) with an overall 14% (9, 21%) increase in energy expenditure in the follow-up test compared to baseline.

#### Resistance Exercise

Eighteen participants (13F) completed pre-intervention baseline and post-intervention follow-up RE acute tests with similar average repetitions, relative intensities for chest press, leg press, and leg extension, and blood lactate concentrations (Ext. Data Figures 6A, 6C, 5B). Total upper body [+44% (30, 86%)], lower body [+28% (18, 39%)], and combined workload [+32% (27, 45%)] increased after the intervention (Ext. Data Figure 6B). Compared to baseline, increases in resistance ranged on average from 18% to 55%, with the largest percent change observed in overhead press and the smallest percent change in leg curl. Absolute weight per set increased for all exercises, with the smallest absolute change occurring in overhead press and the largest in leg press (Supplementary Table 10).

### Biospecimens Collection and Processing

An overview of biospecimen collection success for blood, muscle, and adipose across all time points during the pre-intervention baseline and post-intervention follow-up acute exercise tests is shown in Ext. Data Figure 7 and Supplementary Table 11. Generally, across the time points and tissue type, over 90% success was achieved for blood, muscle, and adipose samples. In total, 31,725 aliquots of processed blood (N=25,236), muscle (N=3,332), and adipose (N=3,157) samples were successfully shipped from the human clinical centers to the centralized biorepository. Further processing was performed at the biorepository with 17,944 samples for blood (10,332), muscle (3,868), and adipose (3,744) sent to various chemical analysis sites for molecular (e.g., genomic, proteomic, metabolomic) analysis^6^.

## DISCUSSION

The pre-COVID phase of the MoTrPAC human clinical trial studied 206 generally healthy sedentary individuals across a wide range of the adult lifespan (18-74 y) and provided insight into the key physiological responses to acute and chronic exercise. The main takeaways from these analyses were 1) the physiological profile (i.e., blood parameters, VO_2_peak, muscular strength) was indicative of a generally healthy and sedentary cohort with typical variation observed across sex and age; 2) the baseline acute exercise bout provided a vigorous exercise challenge for both endurance and resistance exercise with distinct physiological responses between exercise modes; 3) the 12-week progressive endurance and resistance training programs induced robust improvements in several physiological parameters (i.e., VO_2_peak, muscular strength, workload) that varied by exercise mode; and 4) the biospecimen (blood, muscle, and adipose) collections and processing were largely successful across all participating clinical sites. These data highlight the vigorous nature of the acute and chronic exercise procedures, efficacy of the human clinical center MoTrPAC protocol, high fidelity to the protocol, and support molecular interpretations (ref. Molecular Landscape in review at *Nature Medicine*)^13–16^.

The anthropometric and clinical blood biomarkers represented a generally healthy, sedentary cohort^17^. Cardiorespiratory fitness and strength assessments further supported the generally healthy classification when compared to normative values. Cardiorespiratory fitness (e.g., VO_2_peak), is an established prognostic assessment for cardiovascular and all-cause mortality^3,19^. In plots comparing normative values on the cycle ergometer from the FRIEND database of apparently healthy adults ^20^, nearly all (88%) randomized MoTrPAC participants fell within the 10-90^th^ percentiles for cardiorespiratory fitness during their CPET (Figure 3A). As shown by peak RER and RPE at test termination, participants completed a maximal effort to volitional exhaustion. Additional peak values obtained during the CPET, including VE (L·min^−1^), O_2_ pulse (a surrogate for stroke volume, ml•beat^−1^), HR (bpm), BP (mmHg), and workload (watts), were all within ranges for a generally healthy, sedentary population^20–22^, which further supports this classification^17^.

Handgrip strength has gained attention in recent years given its relationship to overall skeletal muscle health, functional status, and prognostic utility for mortality and disability^23,24^. In plots comparing normative handgrip values from the National Institutes of Health (NIH) Toolbox Project^24^, nearly all (84%) randomized MoTrPAC participants fell within the 10-90^th^ percentiles for handgrip strength for their age (Ext. Data Figure 1A). Separate MoTrPAC manuscripts explore cardiorespiratory and strength data associations with molecular associations in blood, muscle, and adipose (ref. Molecular Landscape in review at *Nature Medicine*)^15^. The descriptive profiles suggest sex and age differences align with previous literature^20,22,25–27^, but due to the limited male and older adult representation in the current cohort, these patterns will be explored in the post-suspension cohort (N=1541). Overall, these data coupled with activity screening questionnaires and actigraphy^28,29^ characterize the pre-COVID cohort as generally healthy and sedentary, with typical variances observed across sex and age.

After screening and randomization, participants completed EE, RE, or CON acute bouts, which resulted in distinct responses based on modality. The goal of the EE acute bout was to provide an aerobic steady-state exercise challenge eliciting a robust physiological response known to increase energy demand and perturb numerous molecular networks related to metabolic health^1,2,30–32^. The goal of the RE acute bout was to provide a whole-body exercise challenge comprised of five upper body and three lower body exercises at 10RM intensity to elicit a strong physiological response known to stimulate muscle protein synthesis and the molecular underpinnings for muscle growth^33-39^. As shown in Ext. Data Figure 2, the familiarization-informed workload adjustments were effective in establishing EE and RE acute bout workloads within targeted ranges to support associated molecular interpretations for the prescribed exercise intensities.

As demonstrated by the observed relative intensity (65% VO_2_peak), HRR (70%), RER (0.92), and blood lactate concentrations (4.0 mM), the 40 min EE acute bout was performed at a vigorous intensity and likely near maximal steady-state^40,41^. Relative exercise intensity of 65% of VO_2_peak is considered vigorous for sedentary individuals^42^ and represents ∼4 to 6-fold increase in metabolic rate compared to resting state^43^. As exercise intensity increases, carbohydrates provide a greater contribution to energy demands^32,44–47^. Carbohydrate oxidation, derived from several sources (e.g., blood, liver, and muscle^32,45,48^), increased by ∼10-fold compared to typical resting rates, providing >70% of the energy for the acute exercise bout, while fat oxidation increased by ∼2-fold compared to typical resting values^32,45,48,49^. The increase in metabolic demand and substrate utilization during the vigorous EE acute bout were also apparent at the molecular level in these individuals, including 1) upregulation of glycogen (i.e., storage form of carbohydrate) breakdown pathways in skeletal muscle (ref. Molecular Landscape in review at *Nature Medicine*), 2) upregulation of lipid oxidation and associated metabolites (ref. Molecular Landscape in review at *Nature Medicine*)^14,15^, 3) upregulation of mitochondrial oxidative pathways^13–15^, and 4) upregulation of angiogenic pathways ^13–15^.

The RE acute bout data indicated the whole-body RE dose was vigorous, non-steady-state exercise with a greater reliance on anaerobic metabolism compared to the endurance exercise bout. Acute bout resistances for chest press, leg press, and leg extension elicited intensities aligning with the ACSM estimated 1RM intensity for 10RM (i.e., ∼60-80% 1RM)^50^ and the intensity shown to produce strength increases in novice individuals^51^. Resistance exercise of similar intensity to the MoTrPAC protocol increases protein synthesis^36^, alters mRNA expression^52,53^, greatly alters skeletal muscle metabolism (e.g., ATP turnover, muscle glycogen and triglyceride content)^54,55^, and promotes muscle hypertrophy when repeated over several weeks (i.e., training)^37,39^. Many of these molecular signatures were expanded upon in additional MoTrPAC manuscripts, including 1) upregulation of glycolytic pathways^14^, 2) upregulation of ribosomal biogenesis (ref. Molecular Landscape in review at *Nature Medicine*), and 3) the interplay of protein synthesis and breakdown in skeletal muscle remodeling (ref. Molecular Landscape in review at *Nature Medicine*)^14^.

The EE and RE acute bouts resulted in distinct metabolic perturbations. A notable intermediary of metabolism, lactate, has a rich history of study in exercise physiology^40,56,57^ with its role initially thought to be a waste product^58^. It is now well-established that lactate is a substrate for many tissues (e.g., heart, skeletal muscle, brain, liver) and a diverse signaling molecule^58^ potentially involved in the regulation of transcription factor EB (TFEB) as discussed in the molecular companion manuscript (ref. Molecular Landscape in review at *Nature Medicine*). Blood lactate concentration can rise from ∼0.5-1.0 mM at rest to greater than 20 mM during exercise^59,60^ with submaximal steady-state exercise generally producing a blood lactate between 2-4 mM, which can rise slightly above this value in untrained individuals^40,61^. The MoTrPAC EE participants were on the upper end of this submaximal, steady-state lactate level, which suggests they were likely near maximal steady-state during the exercise bout^40,62^ (Figure 5A-C). The blood lactate following RE acute bout completion was nearly double the steady-state submaximal response observed with the EE acute bout, suggesting a greater anaerobic stimulus to the working muscle^55,59,63–65^.

The goal of the intervention was to provide a vigorous, mode-specific, and progressive exercise training program 3 days per week over 12 weeks followed by a re-evaluation of several phenotypic characteristics and a repeat of the acute exercise test with biospecimen collections (Figure 2C). The improvements in VO_2_peak with the EE^47,66-70^ and RE^71–74^ training programs are in range with previous studies that used a vigorous training stimulus over a similar timeframe. Fewer data are available for VO_2_peak improvements with RE. However, these initial MoTrPAC results are on the upper end of the spectrum, which may be due to the whole-body nature and design of the protocol that will be further examined in the larger cohort. The targeted strength training increases in the RE group were generally observed when compared to EE training and CON^42,72,75,76^. These EE and RE training adaptations were robust for a 12-week vigorous training stimulus and provide strong support for the efficacy of the MoTrPAC exercise training program^42,68,75–79^.

The goal of the post-intervention follow-up EE acute test was to re-evaluate the pre-intervention aerobic steady-state exercise bout at the same relative intensity (65% of VO_2_peak) to compare the physiological response between the untrained and trained states. These preliminary results highlight the adaptation to the MoTrPAC training program and are consistent with previous literature showing increased workload, oxygen uptake, and glucose oxidation at the same relative exercise intensity in the trained state^44,46,80^. The goal of the post-intervention follow-up RE acute bout was to re-evaluate the whole-body exercise at the same relative 10RM intensity to compare the response between the untrained and trained state. The strength increases with the MoTrPAC program were in range with prior studies including young and old individuals with similar vigorous resistance exercise programs^37,72,74–76,81–83^.

The current manuscript aimed to present results from this small, randomized experiment to highlight feasibility and to generate hypotheses for future analyses in the larger cohort of the MoTrPAC study. Results should be interpreted with caution for several reasons: 1) small sample sizes may limit generalizability of findings^84^; 2) simple randomization of small groups can create imbalances in both known and unknown confounders^85,86^; and 3) NIH initiatives have long emphasized caution in interpreting small studies to enhance reproducibility^87^. The main post-suspension MoTrPAC study will include a larger cohort of 1,541 participants randomized under a slightly modified protocol^17^.

Although blocked randomization with site-based stratification was employed, discrepancies in regulatory approvals, start-up times, and pandemic-related interruptions resulted in some imbalance between groups and small sample sizes. Specific limitations in the baseline (pre-intervention) data include: 1) subgroup sample sizes based on intervention group, age group, sex, and time point were as small as four participants. This was particularly noteworthy in the 60+ y age group, so the 40-59 y and 60+ y age groups were combined into one age group (40+ y) for the current manuscript. Additionally, throughout the study, females composed over 70% of the participant cohort; 2) limited generalizability, as 51% of CON participants with biospecimen samples were randomized at two of the ten sites, and 73% of CON participants at four sites; and 3) there was insufficient data to disentangle sex from body composition due to variations in dual-energy x-ray absorptiometry (DXA) machine manufacturers and models across sites, which is being resolved for the larger study.

The pre-COVID phase of the adult sedentary MoTrPAC program highlighted the successful implementation of this complex human clinical trial across all participating centers. The acute and chronic vigorous exercise components showed strong physiological perturbations to the acute exercise challenge and robust adaptations to the training program. The temporal biospecimen (blood, muscle, and adipose) collections and processing that were coupled to the acute exercise bouts were highly successful. The endurance and resistance exercise induced distinct acute and chronic physiological responses, which provided a framework to interrogate the molecular basis for health adaptations to these two popular exercise modalities. These initial data from 206 participants will be expanded upon in the larger post-suspension cohort, allowing for a more comprehensive analysis of the inter-individual variation and adaptation in physiological and molecular exercise responses in humans.

## METHODS

### Participants

A total of 206 generally healthy, sedentary female and male adults were consented, screened, and randomized into intervention groups through block randomization with site-specific stratification. Participant characteristics of the 80 EE, 81 RE, and 45 CON participants are highlighted in Table 1. Due to the COVID-19 suspension, participants completed various phases of the protocol as shown in the CONSORT diagram (Figure 1). Prior to volunteering for this research study, participants were briefed on the project objectives and testing procedures by a member of the investigative team. Participants were informed of the risks and benefits of the research and gave their written consent prior to undergoing study visits. The study was approved by the Institutional Review Board (IRB) of Johns Hopkins University School of Medicine that served as the central IRB for the adult clinical centers for this part of the study (IRB protocol #JHMIRB5; approval date 05/06/2019). This study was conducted in accordance with the Declaration of Helsinki. Volunteers were initially screened as outlined in the MoTrPAC Manuals of Operating Procedures (MOPs) (https://motrpac-data.org/Methods) to ensure they met all eligibility criteria and verify they were healthy and medically cleared to be formally randomized into the study.

### Overall Study Design

The aim of the MoTrPAC study was to recruit sedentary adults across the lifespan who, after screening, were randomized to either a non-exercise CON condition or one of two modes of exercise training: EE or RE. The following sections describe the MoTrPAC protocols utilized in the cohort prior to the COVID-19 suspension. Volunteers were initially screened to ensure they met all eligibility criteria and verify they were healthy and medically cleared to be formally randomized into the study. Participants were then randomized into intervention groups stratified by clinical site. Once randomized, EE and RE participants completed familiarization sessions and a pre-intervention baseline acute endurance or resistance exercise test with biospecimen sampling. CON participants completed biospecimen sampling with 40 min of supine rest in place of an exercise test. Following the pre-intervention baseline acute test, participants began 12 weeks of exercise training or control conditions. Upon completion of the respective interventions, participants repeated phenotypic assessments and the acute test [i.e., post-intervention follow-up testing], including biospecimen sampling. For more detail, please refer to the comprehensive protocol described by MoTrPAC^17^ along with the MOPs (https://motrpac-data.org/Methods) used by clinical sites during data collection.

### CONSORT Diagram

The CONSORT Diagram (Figure 1) reflects randomization and post-randomization activity for the sedentary participants beginning August 2019 through the suspension of activities in March 2020 due to the COVID-19 pandemic. Participant data were broken down by randomized group assignment. The pre-intervention baseline acute test signified the number of participants who had initiated a baseline acute test and had any biospecimens collected.

After pre-intervention baseline acute testing, participants began 12 weeks of EE, RE, or CON intervention. This stage commenced within a median of 11 (7, 13; EE) and 12 (7, 13; RE) days after the acute test. CON intervention timeline started after the baseline acute test. The intervention was originally scheduled for 12 weeks; however, due to the COVID-19 pandemic, participants who completed 8 weeks of the intervention (24 documented sessions) were allowed to complete follow-up assessments and acute testing. Withdrawal from the intervention was defined as a participant who was unwilling or unable to continue with training but was willing to continue with study follow-up assessments.

The same pre-intervention baseline phenotypic assessments were completed during the last 2 weeks of the intervention period. Two to three days after the final intervention session, a post-intervention follow-up acute exercise test was completed, including biospecimen collection. The CONSORT diagram illustrates how many participants provided any biospecimen as part of the acute test.

The number of withdrawals and lost to follow-up (LTFU) between each phase of the study is provided along with LTFU due to COVID-19 suspension. Any participant who stopped attending or was unable to attend the intervention was designated as withdrawn, regardless of the reason. LTFU defines situations where the clinical site was unable to contact the participant after multiple attempts.

### Phenotypic Assessments

Phenotypic assessments included anthropometrics, resting HR and BP, a fasted blood panel, cardiorespiratory fitness, and muscular strength. Body composition was assessed by DXA, but harmonization across different instrument manufacturers and models has not been completed in time for manuscript inclusion. For additional details regarding phenotypic assessments, please refer to the MoTrPAC clinical protocol manuscript^17^ and the MoTrPAC MOPs (https://motrpac-data.org/Methods).

#### Anthropometrics

Each of the measurements was taken in triplicate. Height measurements were obtained using a calibrated stadiometer, weight was measured using an electronic scale, and a Gulick tape measure was used to measure waist circumference at the level of the iliac crest with four ounces of tension. The average of the first two valid measurements was taken, or the average of all three measurements if no two were within the valid range. Valid height measures were expected to be within 0.5 cm, valid weight measures were expected to be within 2 kg, and valid waist circumference measures were expected to be within 1 cm. The average height and average weight were used to calculate body mass index (BMI, kg·m^−2^).

#### Resting Heart Rate and Blood Pressure

Resting HR and BP were measured in triplicate using an automated BP monitor^17^. Resting HR was measured during multiple screening assessments and defined as the lowest value recorded before the pre-intervention baseline acute test. The lowest resting HR measurement taken during post-intervention follow-up phenotypic testing was used as follow-up resting HR.

#### Fasted Blood Panel

Blood samples were collected to assess complete blood count, blood chemistry and metabolic panels, lipid profile, thyroid-stimulating hormone concentration, and hemoglobin A1c (HbA1c) after a ≥10 hr fast (only water allowed).

#### Cardiorespiratory Fitness

Cardiorespiratory fitness was assessed using a CPET on a cycle ergometer (Lode Excalibur Sport Lode BV, Groningen, The Netherlands) with indirect calorimetry. Various ramp protocols were available for selection to accommodate variability in the cardiorespiratory fitness of study participants^17^. HR [monitored by electrocardiogram (ECG)] and BP measurements were obtained prior to, during, immediately post, and during recovery from the CPET. Peak oxygen consumption (VO_2_peak, L·min^−1^ and mL·kg^−1^·min^−1^), VE (L·min^−1^), O_2_ pulse (VO_2_peak / peak HR; mL·beat^−1^), RER, workload (W), and HR (bpm) were collected. All CPETs were required to reach 1.05 RER criteria or were adjudicated to ensure a maximal effort was obtained. VO_2_peak was calculated as the average of the final two completed 30 sec intervals during the CPET. Individual VO_2_peak data were compared to normative values on the cycle ergometer for apparently healthy adult males and females from the Fitness Registry and the Importance of Exercise National Database (FRIEND) (3662 females, 2439 males)^20^ across the lifespan.

#### Muscular Strength

Quadriceps strength was determined by maximal isometric knee extension of the right leg at a 60° knee angle using a suitable strength dynamometer^17^. Grip strength of the dominant hand was obtained using the Jamar Handheld Hydraulic Dynamometer (JLW Instruments, Chicago, IL). Individual handgrip strength was compared to normative values from the National Institutes of Health (NIH) Toolbox Project^24^ comprised of 1232 adults (783 females, 449 males) across the lifespan. Isometric knee extension and grip strength were both measured in triplicate. A fourth attempt was completed for isometric knee extension if the three attempts had >10% coefficient of variation.

#### Accelerometry

Wearable activity monitors (Actigraph GT9X Accelerometer, Pensacola, FL) were used in the MoTrPAC study to objectively assess and quantify the intensity and patterns of free-living activity and sedentary behavior, including sleep. Randomized participants wore the Actigraph GT9X device on their non-dominant wrist at baseline and at approximately the 4th, 8th, and 12th weeks of the intervention. The goal was to obtain about one week of compliant data at each time point, with a compliant day defined as ≥12 hr of wear time. Participants returned the devices after a week of wear, and data were uploaded to Actigraph CentrePoint as previously described^17^. At baseline, Actigraph devices were distributed to participants before randomization. Baseline data collection was typically completed prior to the acute test, with participants removing the device during any familiarization and assessment sessions to capture their normal baseline activity level. Minute-level data (e.g., 3-axis counts, step count) were obtained from Actigraph CentrePoint, with wear time detected using Choi’s algorithm^88^. Daily step counts were averaged over 5 to 8 compliant days at baseline to quantify the usual activity behavior of the participants.

### Endurance Exercise Familiarization Sessions and Acute Bout

Prior to the pre-intervention baseline acute exercise bout and biospecimen collection, EE participants completed two 20 min familiarization sessions on the cycle ergometer (Lode Excalibur Sport Lode BV, Groningen, The Netherlands) separated by ≥48 hr to determine the workload that would elicit ∼65% VO_2_peak. At the end of the intervention, these two familiarization sessions were completed again to establish a post-intervention follow-up acute bout workload based on the follow-up CPET. Follow-up acute bout workload remained at a target of 65% VO_2_peak regardless of intensity obtained during the baseline acute bout.

The pre-intervention baseline acute EE bout occurred 5-8 days after the second familiarization, and the post-intervention follow-up acute EE bout occurred 2-3 days after the last training session. All acute bouts occurred in the morning following ≥10 hr fast. Each acute bout session was composed of three parts: (1) a 5 min warm up at 50% of the estimated power output to elicit 65% VO_2_peak, (2) 40 min cycling at ∼65% VO_2_peak, and (3) a 1 min recovery at ∼25 W (Figure 2A). Biospecimens were collected before, during, and after the completion of the acute EE bout as described in a subsequent section. During the baseline acute bout, a few participants (N=7) stopped prior to completion of exercise due to participant request or safety concerns and experienced symptoms such as feeling faint, fatigue, lightheadedness, and/or dizziness. One EE participant opted not to perform the exercise bout after pre-exercise biospecimen collection. During the follow-up acute bout, 3 EE participants stopped prior to completion of exercise due to feeling faint or other reasons and the workload for one EE participant was decreased to encourage completion.

Respiratory gas exchange data were obtained and averaged over minutes 14:01-17:00 and 34:01-37:00. Substrate utilization [carbohydrate (CHO) and fat (FAT)] was calculated using the following equations based on Jeukendrup et al.^49^ for exercise performed at ∼65% VO_2_peak assuming no protein (urinary nitrogen) contribution:

1. **CHO (g·min^−1^)** = 4.210 (VCO_2_) - 2.962 (VO_2_)
2. **FAT (g·min^−1^)** = 1.695 (VO_2_) - 1.701 (VCO_2_)
3. **CHO (kcal·min^−1^)** = CHO (g·min^−1^) * 4.07 (kcal·g^−1^)
4. **FAT (kcal·min^−1^)** = FAT (g·min^−1^) * 9.75 (kcal·g^−1^)
5. **Energy expenditure (kcal·min^−1^)** = CHO (kcal·min^−1^) + FAT (kcal·min^−1^)
6. **CHO (%)** = CHO (kcal·min^−1^) / energy expenditure (kcal·min^−1^) * 100 Total work and energy expenditure were calculated using the following equations:
7. **Total work (kJ)** = average watts during ∼65% VO_2_peak * ride duration (min) * 60 (sec) / 1000
8. **Total energy expenditure (kcal)** = average energy expenditure (kcal·min^−1^) * ride duration (min)

### Resistance Exercise Familiarization Sessions and Acute Bout

In preparation for the pre-intervention baseline acute exercise bout and biospecimen collection, RE participants completed three familiarization sessions to establish resistances for the five upper body (chest press, overhead press, seated row, triceps extension, biceps curl) and three lower body (leg press, leg curl, leg extension) exercises, with the goal to elicit fatigue within 8-12 repetitions for each of three sets. A 5 min warm-up at 50-60% of HRR was completed on a cycle or treadmill before each familiarization session. The first RE familiarization session was designed to establish individual participant equipment settings, teach proper technique, establish a moderate intensity resistance for each movement, and provide 1-repetition maximum (1RM) practice. The second RE familiarization session required each participant to perform 1RMs for three exercises: leg press, chest press, and leg extension^17^. After the 1RMs were completed, participants performed repetitions, one at a time, for each of the eight exercises to establish a load that elicited a rating of perceived exertion (RPE) of 5-6 on a 0-10 scale (corresponding to a weight eliciting volitional fatigue between 8-12 repetitions). Once this weight was determined, participants performed one set of each exercise to volitional fatigue at this intensity. The third RE familiarization session involved participants performing two sets of each exercise to volitional fatigue at a resistance calculated from data collected during session 2. The weights and repetitions completed during RE familiarization session 3 were used to guide the intensity of the exercises for the RE acute test and to achieve failure at 8-12 repetitions for each set. All familiarization sessions were separated by ≥2 days. The acute RE test occurred 5-8 days after the third RE familiarization session at pre-intervention baseline and 2-3 days after the last RE training session at post-intervention follow-up. All acute tests occurred in the morning following a ≥10 hr fast. After a 5 min warm-up at 50-60% HRR, participants completed three sets each of the five upper and three lower body exercises to volitional fatigue with 90 sec rest between each set (Figure 2A). An additional warm-up set with a resistance ∼70% of first working set was performed for chest press, seated row, and leg press exercises followed by 60 sec rest. At the baseline acute bout, 1 RE participant stopped prior to completion of exercise due to safety concerns. At the follow-up acute bout, all RE participants completed all exercises. Biospecimens were collected before and after the completion of the acute RE bout as described in a subsequent section.

### Control Acute Rest

Participants randomized to CON did not complete exercise familiarization sessions or exercise during their acute tests. CON participants had a period of 15 (11, 24) days between randomization and baseline acute test. Participants rested supine for 40 min to mirror the EE and RE acute test schedule. Biospecimens were collected before, during, and after the completion of the CON rest as described in a subsequent section.

### Acute Test Biospecimen Collection

#### Standardizations for Biospecimen Collections

To standardize conditions prior to the start of each acute test, participants were instructed to 1) refrain from COX-inhibitors for ≥7 days, biotin supplementation for ≥72 hr, and caffeine and alcohol for ≥24 hr, 2) refrain from exercise for 5-8 (EE and RE) or 7-14 (CON) days at pre-intervention baseline or 2-3 days at post-intervention follow-up, 3) and consume a liquid nutritional supplement (Ensure^®^; Abbott Laboratories, Columbus, OH) 10-12 hr before arrival to the clinical center. After ingestion of a liquid nutrition supplement, participants began fasting but were allowed to consume water *ad libitum*. Participants requiring 24 hr biospecimen collections were instructed to continue to follow pre-acute testing instructions and repeat the dietary intake protocol the night before their final biospecimen collection. Participants confirmed compliance with these standardized conditions prior to the acute test.

#### Biospecimen Collection and Processing Procedures

During the acute test, blood, muscle, and adipose samples were collected at specific time points before, during, and after exercise that varied according to each temporal sampling profile (Figure 2B). The post-exercise time point began at the completion of the 40 min cycling, third set of leg extension, or 40 min rest. For CON participants, blood, muscle, and adipose samples were collected during a control rest period that mirrored the timing of the exercise training groups. Upon arrival, participants rested supine for at least 30 min prior to pre-exercise biospecimen collections. After the pre-exercise biospecimen collections, the subsequent collection time points varied for each training group according to their randomized temporal profile. Except for the EE blood collection time points during exercise, participants rested supine for biospecimen collections. Only the baseline time point of each tissue was collected for the post-intervention follow-up CON acute test for all temporal profiles.

The collection and processing protocols for each tissue were previously described in the human clinical protocol design manuscript^17^ and the Biospecimen Collection and Processing MOP (https://motrpac-data.org/Methods). Several criteria were utilized to determine if a biospecimen collection was successfully acquired during each acute test: 1) the collection resulted in at least one processed sample vial; 2) the collection was obtained within the allowable time range for the time point^17^; and 3) the collection provided adequate sample yields required for the study of molecular transducers at the Chemical Analysis Sites (CAS).

Blood lactate was measured as an indicator of exercise intensity at all blood collection time points from baseline through 30 min post-exercise/rest using plasma derived from the EDTA single spin protocol^17^. After plasma aliquot storage at −80°C, lactate concentrations were determined using a Beckman DxC 600 clinical analyzer (Beckman-Coulter, Brea, CA) and LACT 2×50 Reagent (Cat#A95550, Beckman-Coulter, Brea, CA).

#### Intervention Protocols

For all three intervention arms of the study (EE, RE, CON), in-person monitoring visits occurred during weeks 4, 8, and 12, documenting body weight, resting HR, resting BP, self-reported changes in physical activity, dietary patterns, medications, and any safety events related to data collection. If participants reported changes in physical activity or diet, clinical staff counseled participants on the importance of maintaining their lifestyle and remaining weight stable.

#### Endurance Exercise

The EE intervention protocol consisted of 12 weeks of supervised training for 50-60 min per day, 3 days per week, at 60-80% HRR (Figure 2C). MoTrPAC implemented a progressive training program by increasing time (omitting warmup) from 50 min in weeks 1-2 to 60 min for the remainder of the intervention period and increasing intensity from 60% HRR in weeks 1-4, 70% HRR in weeks 5-8, 75% HRR in weeks 9-10, and 80% HRR in weeks 11-12. The intensity was maintained within a range of +/-5% HRR. The exercise time was split evenly between a cycle ergometer and a motorized treadmill (walking or jogging). HR (Zephyr Bioharness BH3, Zephyr Technology Corporation, Annapolis, MD) and RPE (Borg 6-20) were collected during each session. Participants were instructed to maintain their current diet and remain weight stable throughout the duration of the study.

#### Resistance Exercise

The RE intervention protocol consisted of 12 weeks of supervised, progressive resistance exercise training, three days per week (Figure 2C). Each training session lasted approximately 45-60 min, and consisted of five upper body (chest press, overhead press, seated row, triceps extension, biceps curl), and three lower body (leg press, leg curl, leg extension) exercises. Each exercise was performed for 3 working sets at the individual 10RM of each participant, with 90 sec rest between sets. Each session was preceded by a 5 min warm-up on a cycle ergometer at 50-60% HRR. Additionally, the first upper body push-exercise (chest press), the first upper body pull-exercise (seated row), and the first lower body exercise (leg press) had one warmup set (10 reps, 70% 10RM) followed by 60 sec rest preceding the working sets. HR (Zephyr Bioharness BH3, Zephyr Technology Corporation, Annapolis, MD) and RPE (Borg 6-20) were collected during each session. Participants were instructed to maintain diet and remain weight stable throughout the duration of the study.

#### Control

The CON intervention protocol consisted of 12 weeks continuance of sedentary behavior [no more than 1 day per week of regular (structured) endurance or resistance exercise lasting no more than 60 min] (Figure 2C). Participants were asked to maintain their level of physical activity and dietary patterns to remain weight stable. To help promote compliance, clinical staff interacted with CON participants approximately every 2 weeks through adherence calls and in-person monitoring visits. Adherence telephone calls occurred during weeks 2, 6, and 10 to obtain self-reported changes in physical activity and dietary patterns.

#### Quantification and Statistical Analysis

All data shown were summarized as median (25^th^, 75^th^ percentile) for continuous variables and with percentage for categorical variables unless otherwise stated. To summarize outcomes that may have had a variable number of measurements and/or missing responses, mixed models for repeated measures were used. Confidence intervals (95%) were bootstrapped using 1,000 replicates to derive an estimate of uncertainty (Supplementary Table 5). Data analyses were coordinated using R version 4.2.3 with assistance from multiple open-source packages^89–96^.

In general, hypothesis tests and specifically testing for heterogeneity of response in small subgroups were avoided. Our aim was to present results from this small, randomized experiment to highlight feasibility, to generate hypotheses, and to lay a solid foundation for future analyses in the larger cohort of the MoTrPAC study. Aligning with recent guidelines from the American Statistical Association, we emphasize point estimates, measures of dispersion, and confidence limits from hypothesis test^97^. In addition, we defer testing for heterogeneity in subgroups to the larger MoTrPAC cohort, as our analysis was not adequately powered ^98^. As Brookes et al. (Figure 3 of their manuscript) showed^98^, interaction effects must be at least double the main effect to achieve 80% power. While the EE and RE groups each had around 70 participants with biospecimen samples, providing 80% power to detect a 0.5 effect size (difference in means/SD) using a two-sample t-test (alpha = 0.05, two-sided), for tests of interaction effects to have 80% power an interaction effect size > 1 would be required, which is considered large by Cohen^99^. Our aim was to present the results from a hypothesis-generating perspective, following Ioannidis’s cautionary guidance^84^, and to lay a solid foundation for future analyses in the larger cohort of the MoTrPAC study.

## Primary Authors

### Manuscript Writing Group Leads

Anna R. Brandt, Scott Trappe

### Manuscript Writing Group

Jerome Fleg, Bret H. Goodpaster, Byron Jaeger, Christopher A. Jin, Neil M. Johannsen, Daniel H. Katz, Hasmik Keshishian, Wendy M. Kohrt, William E. Kraus, Bridget Lester, Edward L. Melanson, Michael E. Miller, Samuel Montalvo, W. Jack Rejeski, Samiya M. Shimly, Gregory R. Smith, Cynthia L. Stowe

### Lead Analysts

Catherine Gervais, Byron Jaeger, Samuel Montalvo, Courtney Simmons

### Data Generation (*team leads)

*Participant Status:* Cynthia L. Stowe*, Anna R. Brandt, Joseph A. Houmard, Wendy M. Kohrt, W. Jack Rejeski, Courtney Simmons, Michael P. Walkup

*Phenotypic Assessments:* W. Jack Rejeski*, Nicolas Musi*, Anna R. Brandt, Charles F. Burant, Haiying Chen, Jerome Fleg, William E. Kraus, Samuel Montalvo, Cynthia L. Stowe, Jennifer W. Talton

*Endurance Exercise Acute Bout:* Anna R. Brandt*, Brian Coyne, Jerome Fleg, Bret H. Goodpaster, Neil M. Johannsen, William E. Kraus, Edward L. Melanson, Samuel Montalvo, Eric Ravussin, W. Jack Rejeski, Courtney Simmons, Scott Trappe, Todd A. Trappe

*Resistance Exercise Acute Bout:* Neil M. Johannsen*, Haiying Chen, Brian Coyne, William E. Kraus, Edward L. Melanson, Samuel Montalvo, Blake B. Rasmussen, W. Jack Rejeski, Jennifer W. Talton

*Biospecimens:* Bridget Lester*, Paul M. Coen, Shannon S. Emilson, Michael E. Miller, Samuel Montalvo, Jessica L. Rooney, Lauren M. Sparks, Cynthia L. Stowe, Jennifer W. Talton, Scott Trappe

*Endurance Exercise Intervention:* Neil M. Johannsen*, Brian Coyne, Daniel H. Katz, Edward L. Melanson, Samuel Montalvo, W. Jack Rejeski, Michael P. Walkup

*Resistance Exercise Intervention:* Neil M. Johannsen*, Brian Coyne, Daniel H. Katz, Edward L. Melanson, Samuel Montalvo, Blake B. Rasmussen, W. Jack Rejeski, Michael P. Walkup

*Intervention Monitoring:* Anna R. Brandt*, Raymond Jones, Samuel Montalvo, W. Jack Rejeski, Cynthia L. Stowe, Taylor Taylor, Michael P. Walkup

*Accelerometry:* Samiya M. Shimly*, Shyh-Huei Chen, Jason Fanning, John M. Jakicic, Samuel Montalvo, Matthew T. Wheeler

*Analytical Bloods:* Leanna M. Ross*, Byron Jaeger*, William E. Kraus, Michael E. Miller, Michael Muehlbauer, Christopher B. Newgard, Courtney Simmons

### Figure and Table Development

Anna R. Brandt, Catherine Gervais, Byron Jaeger, Michael E. Miller, Samuel Montalvo, Courtney Simmons, Scott Trappe

### Human Clinical Adult Study Leadership

Marcas M. Bamman, Thomas W. Buford, Bret H. Goodpaster, Joseph A. Houmard, John M. Jakicic, Wendy M. Kohrt, William E. Kraus, Nicolas Musi, Blake B. Rasmussen, Eric Ravussin, Scott W. Trappe

### Senior Manuscript Leadership

Bret H. Goodpaster, Neil M. Johannsen, Daniel H. Katz, Edward L. Melanson, Michael E. Miller, W. Jack Rejeski, Scott Trappe

### Lead Contact

Scott Trappe

## The MoTrPAC Study Group

### Consortium Coordination

*Bioinformatics Center - Stanford University, Stanford, CA*
David Amar; Euan Ashley; Jeffrey W. Christle; David Jimenez-Morales; Daniel H. Katz; Malene E. Lindholm; Samuel Montalvo; Jonathan N. Myers; Samiya M. Shimly; Matthew T. Wheeler;

*Biospecimens Repository - University of Vermont, Burlington, VT*
Jessica L. Rooney; Russell Tracy;

*Data Management, Analysis, and Quality Control Center - Wake Forest University School of Medicine, Winston-Salem, NC*
Haiying Chen; Shyh-Huei Chen; Shannon S. Emilson; Catherine Gervais; Fang-Chi Hsu; Michael E. Miller; Eric W. Reynolds; Joseph Rigdon; Courtney Simmons; Cynthia L. Stowe; Jennifer W. Talton; Michael P. Walkup;

*Exercise Intervention Core - Wake Forest University, Winston-Salem, NC*
Jason Fanning; W. Jack Rejeski;

*National Institutes of Health, Bethesda, MD*
Jerome Fleg;

### Chemical Analysis Sites

*Broad Institute of MIT & Harvard (Carr Lab), Cambridge, MA*
Steven Carr; Natalie Clark; Patrick Hart; Hasmik Keshishian;

*Broad (Gerzten Lab), Beth Israel Deaconess Medical Center, Boston, MA*
Robert E. Gerszten; Jeremy M. Robbins;

*Broad (Newgard Lab), Duke University, Durham, NC*
Michael Muehlbauer; Christopher B. Newgard;

*Icahn School of Medicine at Mount Sinai, New York City, NY*
Stuart C. Sealfon; Gregory R. Smith; Yifei Sun; Martin J. Walsh;

*Stanford University, Stanford, CA*
Christopher A. Jin; Michael P. Snyder;

*University of Michigan, Ann Arbor, MI*
Charles F. Burant

### Clinical Sites

*Ball State University, Muncie, IN*
Alicia Belangee; Gerard Boyd; Anna R. Brandt; Toby L. Chambers; Clarisa Chavez; Alex Claiborne; Matthew Douglass; Will Fountain; Aaron Gouw; Bruce Graham; Kevin Gries; Ryan Hughes; Andrew Jones; Dillon Kuszmaul; Bridget Lester; Colleen Lynch; Kiril Minchev; Cristhian Montenegro; Masatoshi Naruse; Ulrika Raue; Ethan Robbins; Kaitlyn Rogers; Chad Skiles; Andrew Stroh; Scott Trappe; Todd A. Trappe; Caroline S. Vincenty; Gilhyeon Yoon;

*AdventHealth Translational Research Institute, Orlando, FL*
Elvis A. Carnero; Paul M. Coen; Bret H. Goodpaster; Lauren M. Sparks;

*The University of Alabama at Birmingham - Birmingham, AL*
Marcas M. Bamman; Thomas W. Buford; Gary R. Cutter; Raymond Jones; Taylor Taylor; Anna Thalacker-Mercer;

*Cedars-Sinai Medical Center, Los Angeles, California*
Arianne Aslamy; Samuel Cohen; Sara Espinoza; Nicolas Musi; Gary Weaver;

*East Carolina University, Greenville, NC*
Joseph A. Houmard;

*Duke University, Durham, NC*
Hiba AbouAssi; Will Bennett; Katherine A. Collins-Bennett; Brian Coyne; Kim M. Huffman; Johanna L. Johnson; William E. Kraus; Leanna M. Ross;

*Pennington Biomedical Research Center, Baton Rouge, LA*
Melissa Harris; Neil M. Johannsen; Adam Lowe; Tuomo Rankinen; Eric Ravussin;

*University of California, Irvine, CA*
Dan M. Cooper; Fadia Haddad; Shlomit Radom-Aizik;

*University of Colorado, Denver, CO*
Nicole Adams; Bryan C. Bergman; Daniel H. Bessesen; Zachary Clayton; Andrew Hepler; Catherine M. Jankowski; Wendy M. Kohrt; Edward L. Melanson; Kerrie L. Moreau; Irene E. Schauer; Robert S. Schwartz; Kristen Sutton;

*University of Pittsburgh, Pittsburgh, PA*
John M. Jakicic; Renee Rogers;

*University of Texas Health Science Center, San Antonio, TX*
Tiffany Cortes; Blake B. Rasmussen; Elena Volpi;

### Preclinical Animal Study Sites

*Joslin Diabetes Center, Boston, MA*
Laurie J. Goodyear;

*University of Iowa, Iowa City, IA*
Sue C. Bodine;

*Virginia Polytechnic Institute and State University, Charlottesville, VA*
Zhen Yan;

## MoTrPAC Study Group Acknowledgements

### Consortium Coordination

*Administrative Coordinating Center - University of Florida, Gainesville, FL*
Ching-ju Lu;

*Biospecimens Repository - University of Vermont, Burlington, VT*
Elaine Cornell; Nicole Gagne; Sandra T. May;

*Data Management, Analysis, and Quality Control Center - Wake Forest University School of Medicine, Winston-Salem, NC*
Andrea Anderson; Jerry Barnes; Kate Boyer; Sarah Bruschi; Brandon Bukas; Haiying Chen; Shyh-Huei Chen; Shannon S. Emilson; Catherine Gervais; Leora Henkin; Fang- Chi Hsu; Byron Jaeger; Michael E. Miller; John Nichols; June Pierce; David Popoli; Eric W. Reynolds; Joseph Rigdon; Jeremy Rogers; Scott Rushing; Santiago Saldana; Courtney Simmons; Debbie Steinberg; Cynthia L. Stowe; Jennifer W. Talton; Michael P. Walkup; Christopher Webb; Sawyer Welden; Jack White; Marilyn Williams; Sharon Wilmoth;

*Exercise Intervention Core - Wake Forest University, Winston-Salem, NC*
Barbara Nicklas;

*National Institutes of Health, Bethesda, MD*
Amanda Boyce; Emily Carifi; Jonelle K. Drugan; Jerome Fleg; Stephanie George; Lyndon Joseph; Padma Maruvada; Concepcion R. Nierras; George Papanicolaou; John P. Williams; Ashley Xia;

### Chemical Analysis Sites

*Broad (Gerzten Lab), Beth Israel Deaconess Medical Center, Boston, MA*
Robert E. Gerszten;

*Georgia Institute of Technology, Atlanta, GA*
Carter Asef; Facundo M. Fernández; David A. Gaul; Samuel G. Moore;

*Icahn School of Medicine at Mount Sinai, New York City, NY*
Megan Januska; Martin J. Walsh;

*Pacific Northwest National Laboratory, Richland, WA*
Joshua N. Adkins; Marina A. Gritsenko; Paul D. Piehowski; Wei-Jun Qian;

*Stanford University, Stanford, CA*
Stephen B. Montgomery; Kevin S. Smith; Nikolai G. Vetr;

*University of Michigan, Ann Arbor, MI*
Charles R. Evans; Maureen T. Kachman; Jun Z. Li;

### Clinical Sites

*Ball State University, Muncie, IN*
Gary Lee;

*Cedars-Sinai Medical Center, Los Angeles, California*
Arianne Aslamy; Kuan Tsen Chen; Susan Cheng; Yunhee Choi-Kuaea; Samuel Cohen; Alice Conde; Gavin Connolly; Fadi Dawood; Sara Espinoza; Sandra Gomez; Isabel Homberg Reissmeier; Shao-Jung Hu; Patricia Iluore; Morvarid Kabir; Nour-Lynn Mouallem; Nicolas Musi; Nina Nguimzon; Stacie Saldin; Tararinsey Seng; Purnima Stanek; Ali Tazhibi; Nevyana Todorova; Corey Tolson; Gary Weaver; Ning Zhang;

*East Carolina University, Greenville, NC*
Patricia Brophy; Nicholas T. Broskey; Gabriel Dubis; Elizabeth Gates; Dominique Jones;

*Duke University, Durham, NC*
Brian Andonian; Cherrie Barnes; Kelsey Belski; Liezl Mae Fos; Andrew Hoselton; Leslie Kelly; Kaileigh Moertl; Megan Reaves; Alyssa Sudnick; Leslie Willis;

*Pennington Biomedical Research Center, Baton Rouge, LA*
Carlante Emerson; Kishore M. Gadde; Chelsea Hendrick; Juan Lertora; Alyssa McCarron; Ryan Nash; Phillip Nauta; Robert L. Newton Jr.;

*University of Colorado, Denver, CO*
Teylar Adelsberger; Caitlin Allison; Gabriel Buxo; Zach Buxo; David Calabrese; Ryan Cloud; Andrew Espinoza; Ellie Gibbons; Laurel Gustafson; Jere’ Hamilton; Nick Jeffers; Brandon Kassel; Vincent Khuu; Jenelynn Kimble; Adrian Loubriel; Kate Meadows; Lucas Medsker; Kelly Monaghan; Omid Nabavizadeh; Claire Newman; Adam Orynczak; Paige (Noella) Patterson; Ruby Pressl; Kathleen Resman; Ellen Rohan; Jennyleigh Santa Rosa; Prajakta Shanbhag; Paige Simonavice; Jacklyn (Jackie) Snedeger; Samantha Sumner; Tracy Swibas; Christy Tebsherani; Rhett Turner; Ryan Wendell; Terry Witten;

*University of Pittsburgh, Pittsburgh, PA*
Susan Barr; Dan Forman; Erin Kershaw; Anne Newman; Brad Nindl; Maja Stefanovic- Racic;

*University of Texas Health Science Center, San Antonio, TX*
Allison Stepanenko;

### Preclinical Animal Study Sites

*Joslin Diabetes Center, Boston, MA*
Michael F. Hirshman;

*University of Florida, Gainesville, FL*
Karyn A. Esser; Mark Viggars;

## DATA AVAILABILITY

MoTrPAC Protocol and Manuals of Operating Procedures (MOPs) are publicly available via https://motrpac-data.org/methods. Data access inquiries should be sent to. MoTrPAC data are publicly available via https://motrpac-data.org/data-access. Data access inquiries should be sent to. The MotrpacHumanPreSuspensionData R package (https://github.com/MoTrPAC/MotrpacHumanPreSuspensionData) contains data objects that correspond to the datasets as described in the methods. The motrpac-precovid-adult- sed-clinic repository (https://github.com/MoTrPAC/motrpac-precovid-adult-sed-clinic) contains code to generate visualizations, tables, and statistics presented in the text, utilizing the MotrpacHumanPreSuspensionData package.

## ADDITIONAL RESOURCES

Please refer to the MoTrPAC data repository website (https://motrpac-data.org/) for datasets and Manual of Operations (MOPS, https://motrpac-data.org/Methods).

## ACKNOWLEDGEMENTS

Authors thank the participants who volunteered and gave time for this study. Graphical abstract created in BioRender. Douglass, M. (2026) https://BioRender.com/9m86gx0.

## FUNDING

The MoTrPAC Study is supported by NIH grants U24OD026629 (Bioinformatics Center), U24DK112349, U24DK112342, U24DK112340, U24DK112341, U24DK112326, U24DK112331, U24DK112348 (Chemical Analysis Sites), U01AR071133, U01AR071130, U01AR071124, U01AR071128, U01AR071150, U01AR071160, U01AR071158 (Clinical Centers), U24AR071113 (Consortium Coordinating Center), U01AG055133, U01AG055137 and U01AG055135 (PASS/Animal Sites). Additional funding supporting this study include: J.M.R: K23 HL150327, R03OD038387; D.H.K: K23HL164980, 23CDA1040581; B.R: 5U01AR071150, 2U01AR071130; E.R: P30DK072476; K.C.B: 1K01HL177266-01A1; K.M.: R01AG089069; N.M and S.E: P30AG094848; Z.C: NIH R00 HL159241; R.G: R01NR019628, R01DK081572, 21CVD01 (Leducq Foundation), R01HL133870; M.E.L, S.M, and M.T.W: The Wu Tsai Human Performance Alliance at Stanford and the Joe and Clara Tsai Foundation; T.C. and E.V: P30AG044271.

## DISCLOSURES

B.H.G has served as a member of scientific advisory boards; J.M.R is a consultant for Edwards Lifesciences; Abbott Laboratories; Janssen Pharmaceuticals; M.P.S is a cofounder and shareholder of January AI; S.A.C is on the scientific advisory boards of PrognomIQ, MOBILion Systems, Kymera, and Stand Up2 Cancer; S.C.S is a founder of GNOMX Corp, leads its scientific advisory board and serves as its temporary Chief Scientific Officer; E.A.A is: Founder: Personalis, Deepcell, Svexa, Saturnus Bio, Swift Bio. Founder Advisor: Candela, Parameter Health. Advisor: Pacific Biosciences. Non- executive director: AstraZeneca, Dexcom. Publicly traded stock: Personalis, Pacific Biosciences, AstraZeneca. Collaborative support in kind: Illumina, Pacific Biosciences, Oxford Nanopore, Cache, Cellsonics; G.R.C is a part of: Data and Safety Monitoring Boards: Applied Therapeutics, AI therapeutics, Amgen-NMO peds, AMO Pharma, Argenx, Astra-Zeneca, Bristol Meyers Squibb, CSL Behring, DiamedicaTherapeutics, Horizon Pharmaceuticals, Immunic, Inhrbx-sanfofi, Karuna Therapeutics, Kezar Life Sciences, Medtronic, Merck, Meiji Seika Pharma, Mitsubishi Tanabe Pharma Holdings, Prothena Biosciences, Novartis, Pipeline Therapeutics (Contineum), Regeneron, Sanofi-Aventis, Teva Pharmaceuticals, United BioSource LLC, University of Texas Southwestern, Zenas Biopharmaceuticals. Consulting or Advisory Boards: Alexion, Antisense Therapeutics/Percheron, Avotres, Biogen, Clene Nanomedicine, Clinical Trial Solutions LLC, Endra Life Sciences, Genzyme, Genentech, Immunic, Klein-Buendel Incorporated, Kyverna Therapeutics, Inc., Linical, Merck/Serono, Noema, Neurogenesis, Perception Neurosciences, Protalix Biotherapeutics, Regeneron, Revelstone Consulting, Roche, Sapience Therapeutics, Tenmile. G.R.C is employed by the University of Alabama at Birmingham and President of Pythagoras, Inc. a private consulting company located in Birmingham AL. J.M.J is on the Scientific Advisory Board for Wondr Health, Inc.; P.M.J.B is currently an employee at Pfizer, Inc., unrelated to this project; R.R is a scientific advisor to AstraZeneca, Neurocrine Biosciences, and the American Council on Exercise, and a consultant to Wonder Health, Inc. and seca.; B.J receives consulting fees from Perisphere Real World Evidence, LLC, unrelated to this project.

## DISCLAIMERS

The content of this manuscript is solely the responsibility of the authors and does not necessarily represent the views of the National Heart, Lung, and Blood Institute, the National Institutes of Health, or the United States Department of Health and Human Services.

## AUTHOR CONTRIBUTIONS

All authors reviewed and revised the manuscript. Detailed author contributions are provided in the Supplementary Information.

## Extended Data Figures

**Extended Data Figure 1.**
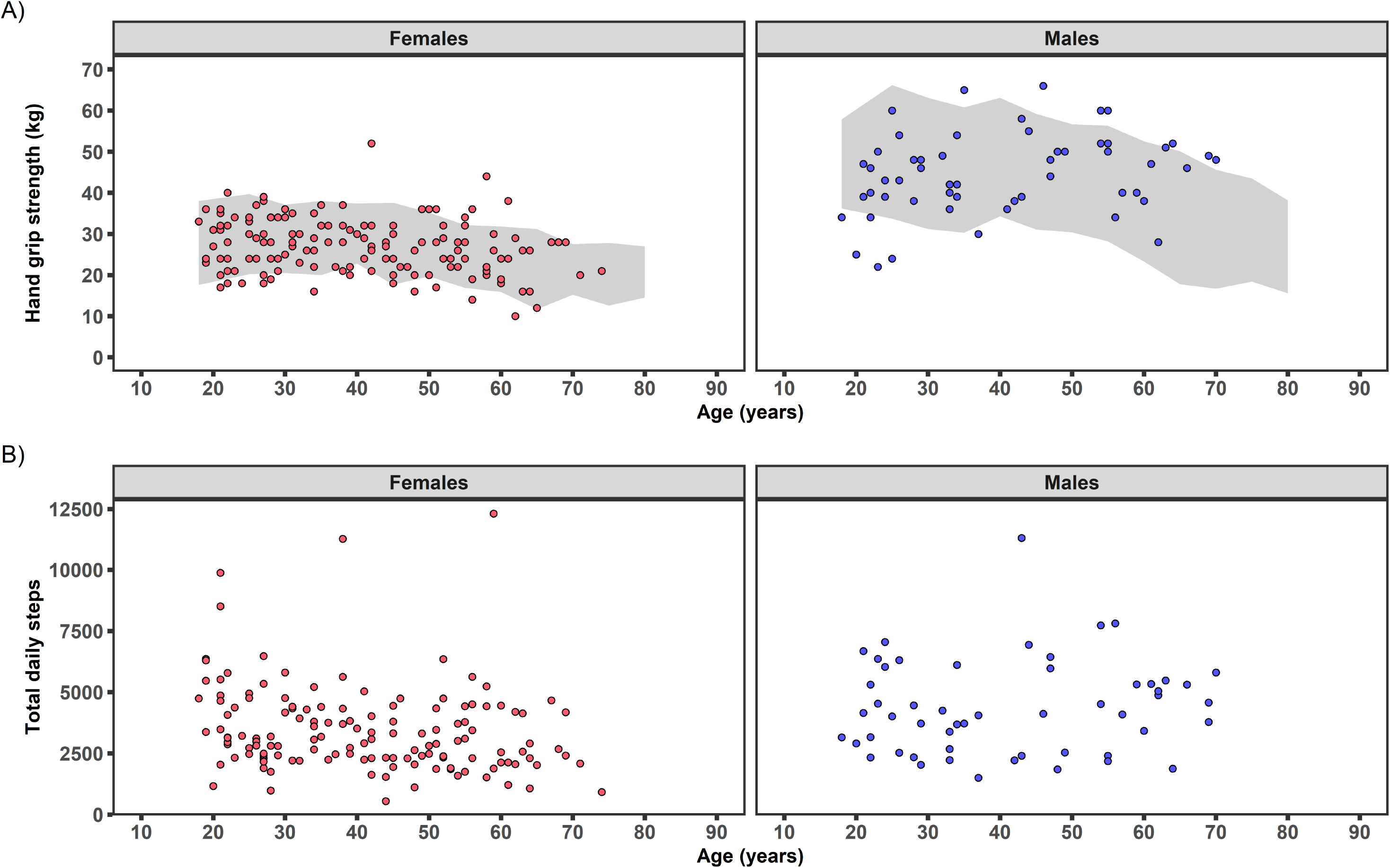
Individual handgrip strength and daily step count per day graphed by age. A) Individual maximal handgrip strength at baseline for randomized participants (N=206; 149F, 57M) are graphed by age and compared with normative values from the NIH Toolbox Project (783 females, 449 males)^24^. Grey area represents 10^th^-90^th^ percentiles of NIH Toolbox Project database across the adult lifespan. B) Individual average step count per day for females (N=140) and males (N=53) were graphed by age. Data were calculated from average of 5-8 compliant days during baseline Actigraph collection.

**Extended Data Figure 2.**
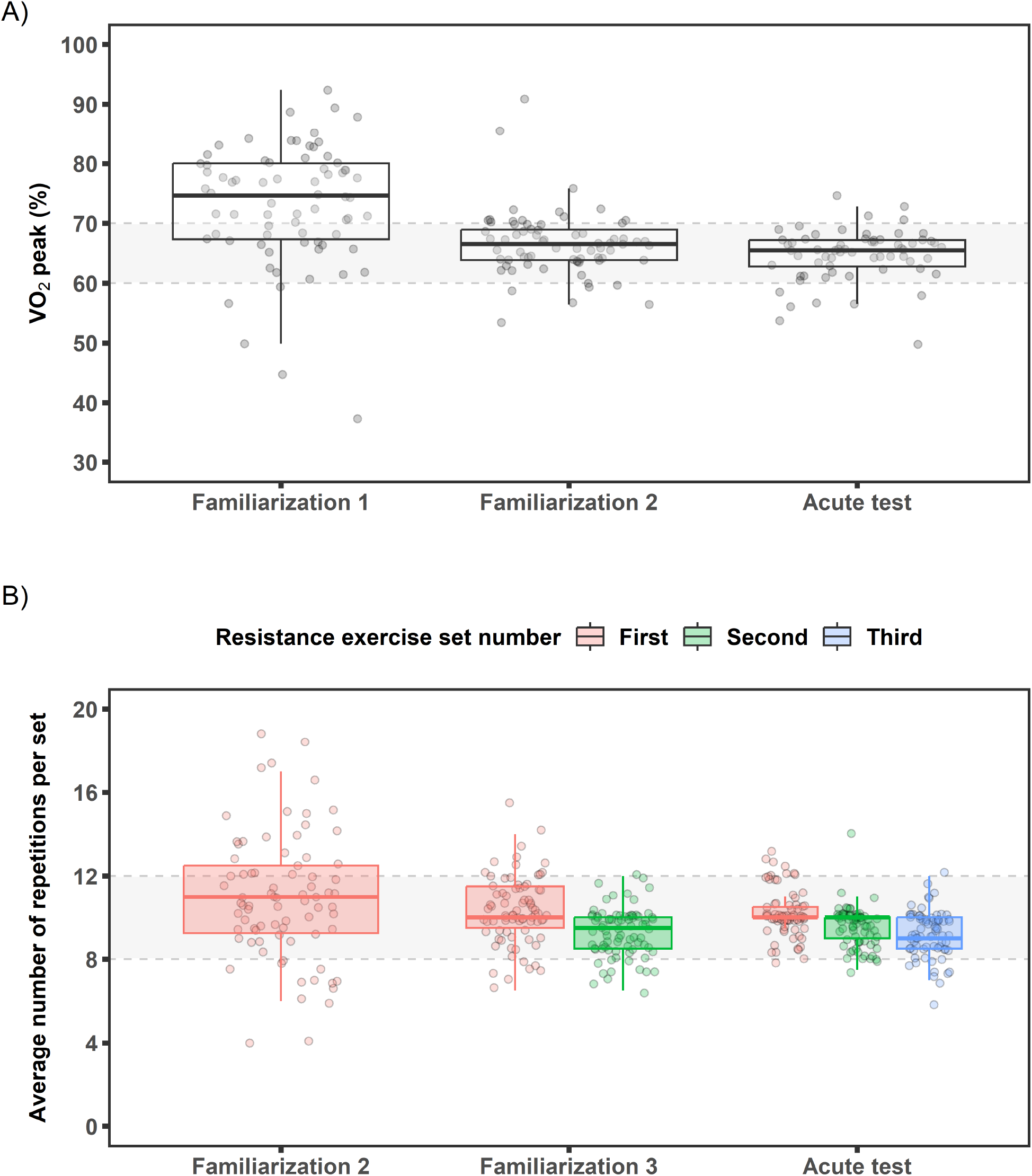
Baseline endurance and resistance familiarization sessions and acute bout exercise intensities. A) Endurance exercise intensity (percent VO_2_peak) is plotted during familiarization sessions and acute bout. Average VO_2_ (L·min^−1^) values for the last two minutes of the familiarization session are plotted. Acute bout values are from averaging minutes 14-17 with 34-37 of exercise. Goal intensity of 65±5% VO_2_peak is shown in grey by the dashed lines. B) Resistance exercise intensity (average number of maximal repetitions per set) completed at familiarization sessions 2 and 3 and acute bout. Participants completed exercises to failure at these visits with 1 set at familiarization 2, 2 sets at familiarization 3, and 3 sets at acute bout. Dashed lines and grey shading show target range of 8-12 repetitions for 10RM. Familiarization session 1 was primarily an introduction to equipment and technique, thus not included. Data shown are boxplots with individual observations.

**Extended Data Figure 3.**
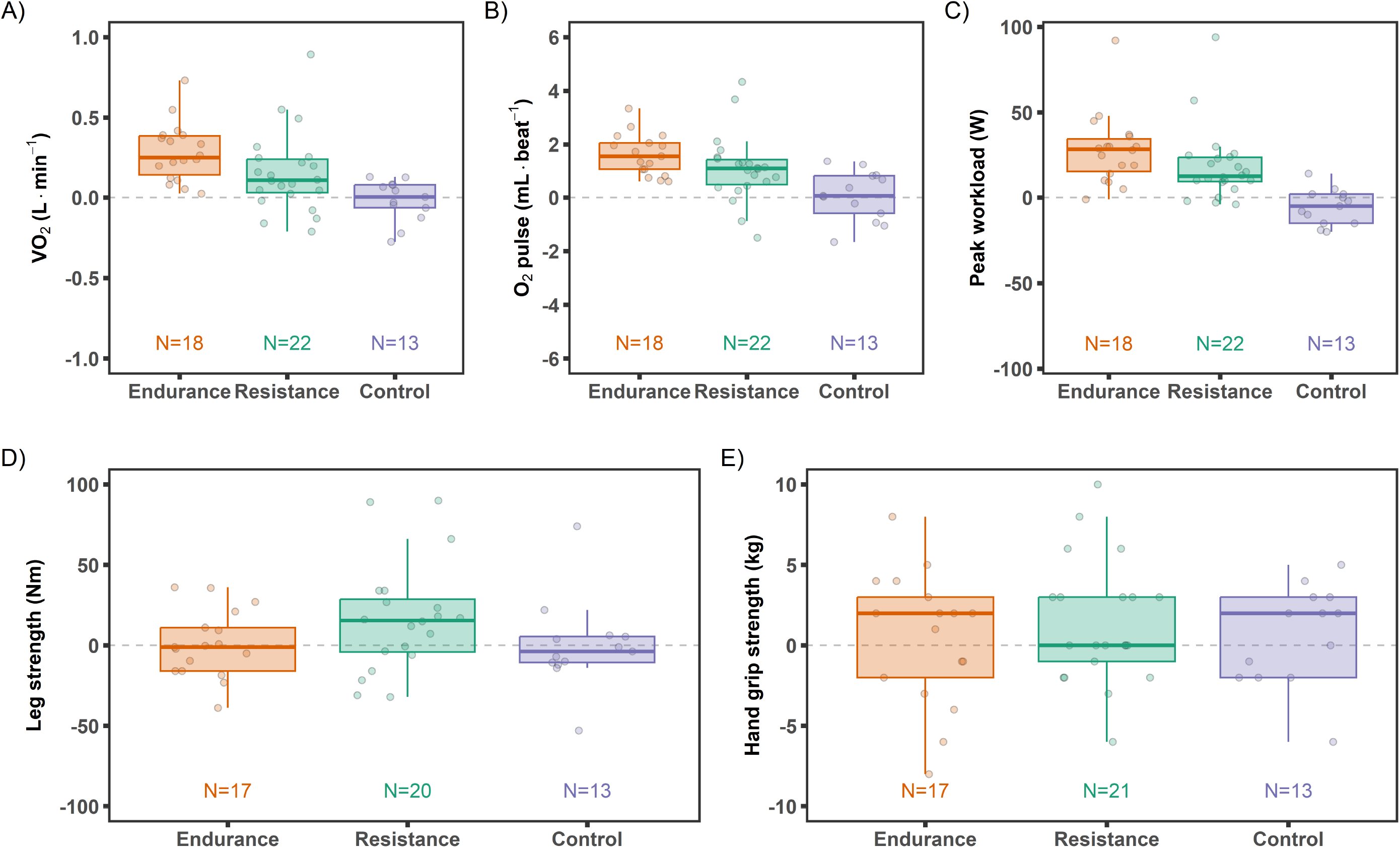
Absolute change in maximal CPET and strength metrics from pre-intervention baseline to post-intervention follow-up. Change in VO_2_peak (L·min^−1^) (A), peak O_2_ pulse (mL·beat^−1^) (B), and peak workload (W) (C) from the cardiopulmonary exercise test (CPET). Change in strength measurements including maximal leg strength [isometric knee extensor strength (Nm)] (D) and maximal hand grip strength (kg) (E).

**Extended Data Figure 4.**
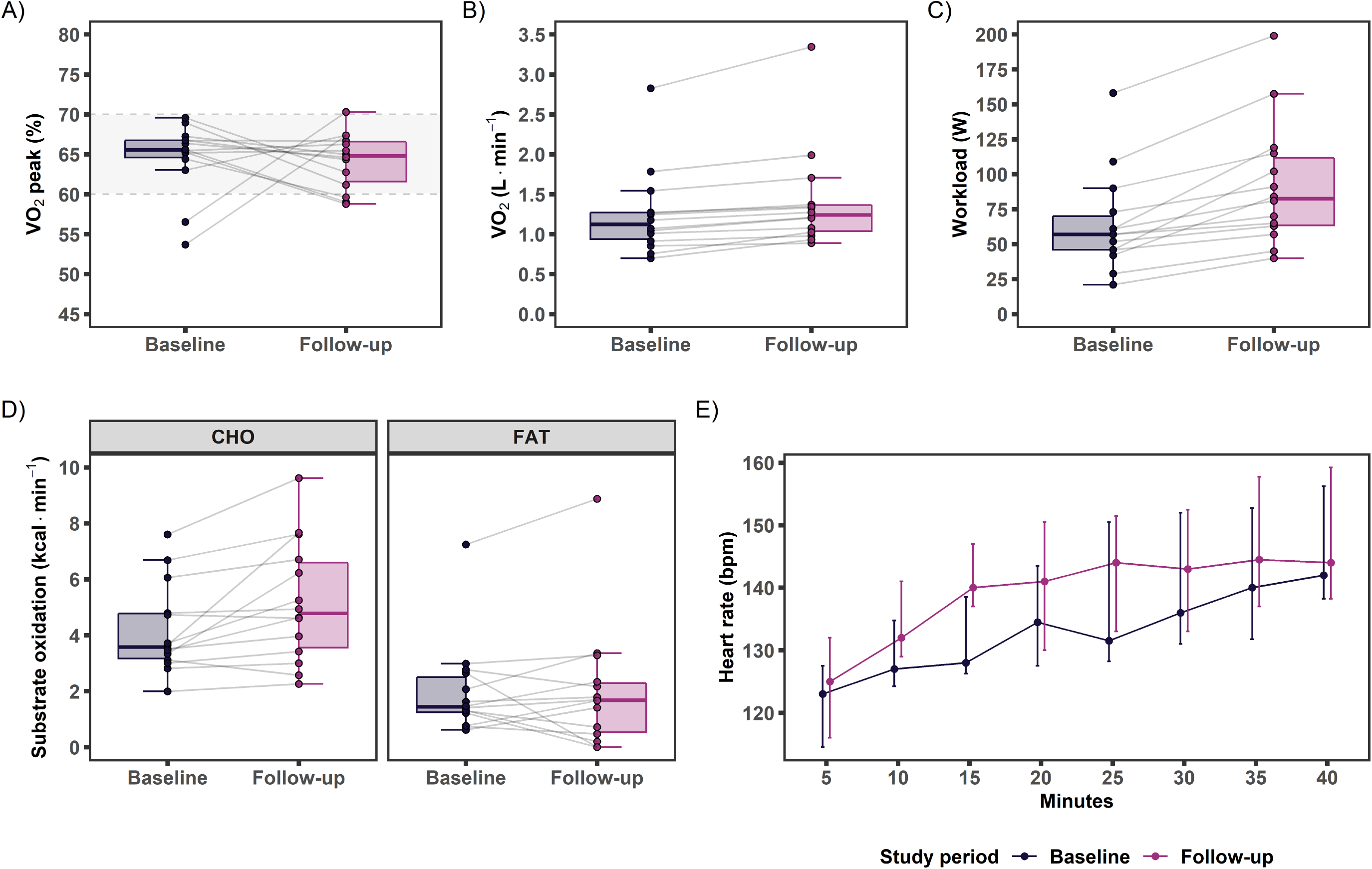
Pre-intervention baseline and post-intervention follow-up endurance exercise acute bout parameters. A-C) Exercise intensity shown as (A) percent VO_2_peak, (B) oxygen uptake (L·min^−1^), and (C) workload (W). Target exercise intensity range of 65±5% VO_2_peak is indicated by the dotted lines and grey shading in panel A. D) Average carbohydrate (CHO) and fat utilization during the acute test. E) Following the 5 min warm-up, the median (25^th^-75^th^ percentile) heart rate progression is plotted during the 40 min acute bout over 5 min intervals. Data shown in panels A-D are boxplots and individual observations. Individual observations in panels A-D are 6 min averages of minutes 14-17 with 34-37.

**Extended Data Figure 5.**
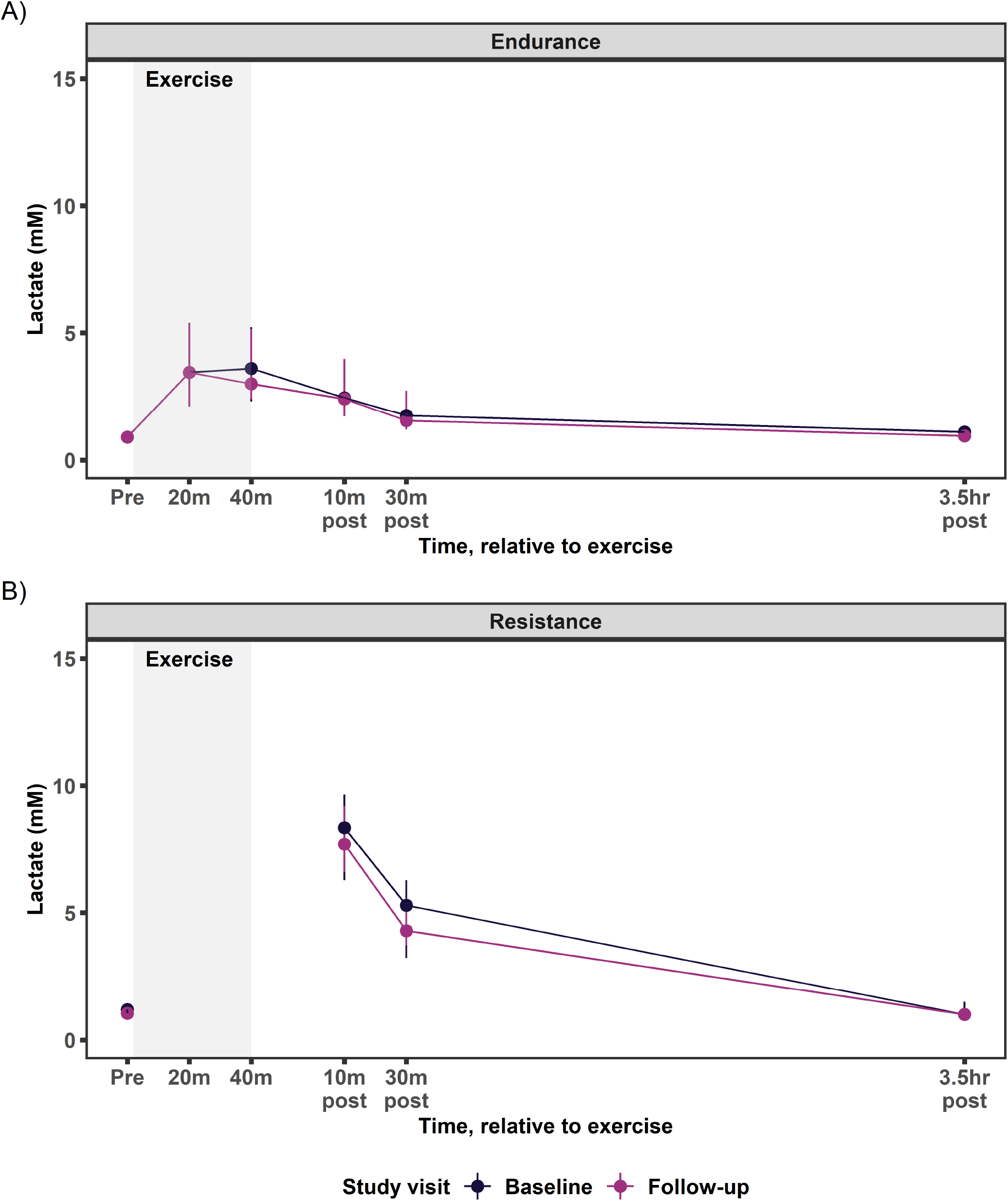
Venous blood lactate from pre-intervention baseline and post-intervention follow-up acute endurance and resistance tests. Pre-intervention baseline and post-intervention follow-up endurance (A) and resistance (B) acute test blood lactate. Data shown as median (25^th^-75^th^ percentile).

**Extended Data Figure 6.**
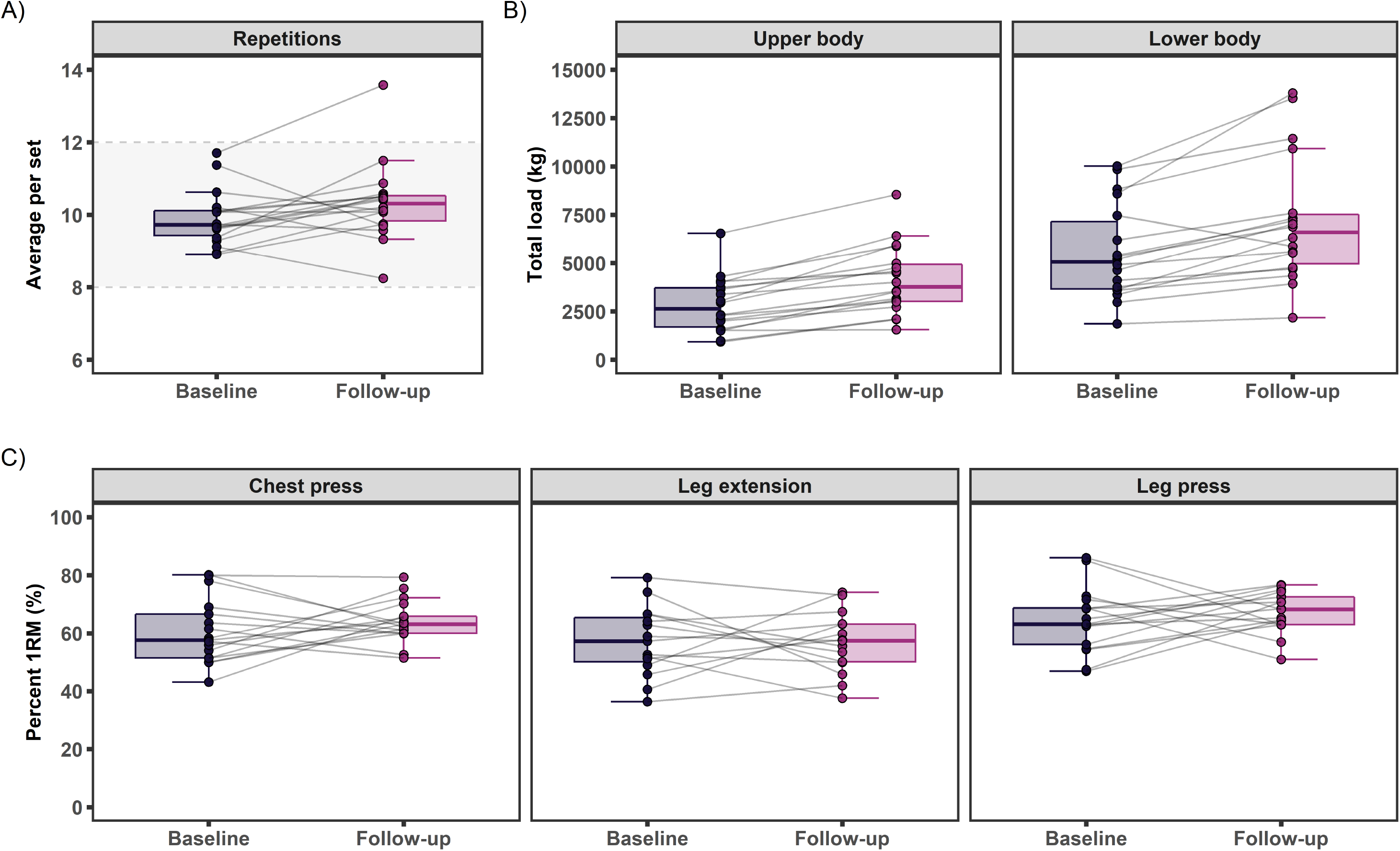
Pre-intervention baseline and post-intervention follow-up resistance exercise acute bout parameters. A) Average number of repetitions per set on all working sets plotted with the 8-12 repetition goal, which is indicated by the dotted lines and shading. B) Total load [weight (kg)*repetitions*sets] for lower and upper body. Total load was plotted for participants that completed at least 1 exercise. C) Average percent one repetition maximum (%1RM) for chest press, leg press, and leg extension. Data shown are boxplots with individual observations drawn as points.

**Extended Data Figure 7.**
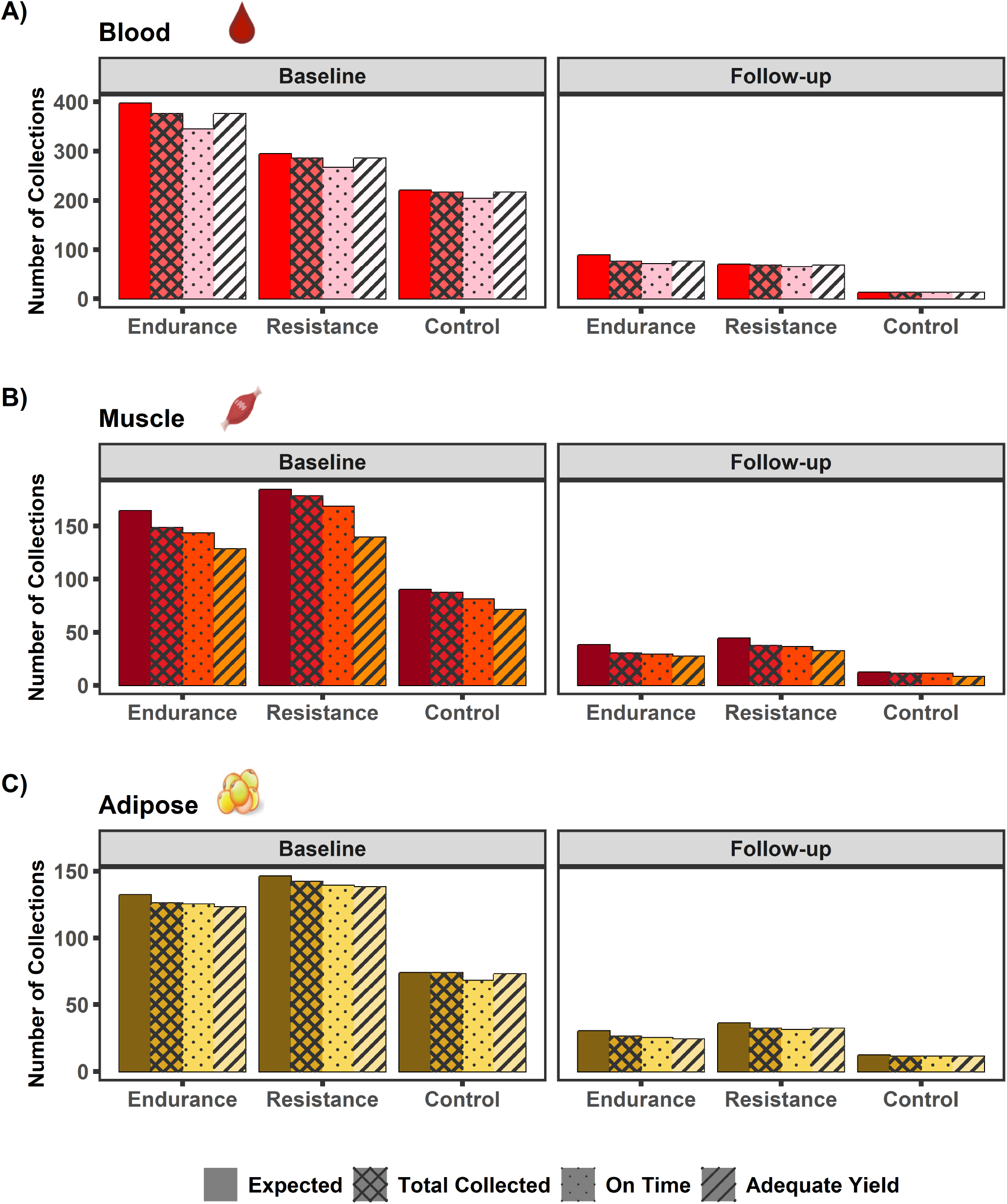
Biospecimen collection success. Baseline acute test and follow-up biospecimen collections for (A) blood, (B) skeletal muscle, and (C) adipose. Fewer biospecimen samples were expected at follow-up due to the COVID-19 suspension. For blood collections, only EDTA Single Spin Plasma and PAXGene RNA sample types are included since these were collected at each sampling time point for all temporal profiles and were involved in the main chemical analyses. Expected = Number of expected collections for acute tests performed. Total Collected = Number of collections that resulted in at least one processed sample vial. On Time = Number of collections that resulted in at least one processed sample vial and was collected within the allowable time range for the time point. Adequate Yield = Number of collections that resulted in adequate yield for shipment to CAS platform.

## SUPPLEMENTARY TABLES FOUND IN EXCEL FILE

**Supplementary Table 1.** Pre-intervention baseline characteristics of participants by sex and age group.

**Supplementary Table 2.** Pre-intervention baseline characteristics of participants who initiated the baseline acute test by randomized intervention group.

**Supplementary Table 3.** Pre-intervention baseline characteristics of participants by baseline acute test initiation status.

**Supplementary Table 4.** Pre-intervention baseline characteristics of participants with post-intervention follow-up data by randomized intervention group.

**Supplementary Table 5.** Pre-intervention baseline endurance exercise acute bout parameters by sex and age group.

**Supplementary Table 6.** Pre-intervention baseline resistance exercise acute bout parameters by age and sex.

**Supplementary Table 7.** Exercise intervention data summary by week.

**Supplementary Table 8.** Baseline and follow-up characteristics of participants with follow-up phenotypic data by randomized intervention group.

**Supplementary Table 9.** Baseline and follow-up endurance exercise acute bout parameters.

**Supplementary Table 10.** Baseline and follow-up resistance exercise acute bout parameters.

**Supplementary Table 11.** Overview of biospecimen collection success for each sample type at baseline and follow-up.

## Notes

https://motrpac-data.org/

### Summary of Updates

Manuscript text has been reorganized with minor updates to content. Figures and tables have also been reorganized with a core figure/table package in the main manuscript and additional items added to supplemental/extended data files.

